# Phenotype switching of the mutation rate as a strategy for crossing fitness valleys

**DOI:** 10.1101/2021.07.14.452333

**Authors:** Gabriela Lobinska, Yitzhak Pilpel, Yoav Ram

## Abstract

Mutation rate plays an important role in adaptive evolution due to its effect on the rate of appearance of both beneficial and deleterious mutations and is therefore subject to second-order selection. The mutation rate varies between and within species and populations, increases under some stresses, and can be modified by mutator and anti-mutator alleles. It may also vary among genetically identical individuals: empirical evidence from bacteria suggests that the mutation rate can be affected by translation errors and expression noise in various proteins. Importantly, this non-genetic variation may be heritable via a transgenerational epigenetic mode of inheritance, giving rise to mutator phenotype switching that is independent from mutator alleles. Here we investigate mathematically how the rate of adaptive evolution on rugged, complex fitness landscapes is affected by the rate of mutation rate phenotype switching. Motivated by recent experimental results of mutation rate variation, we model an asexual population with two mutation rate phenotypes, non-mutator and mutator. An offspring may switch from its parental phenotype to the other phenotype. Thus, the mutation rate can be interpreted as a genetically inherited trait when the switching rate is low, as an epigenetically inherited trait when the switching rate is intermediate, or as a randomly determined trait when the switching rate is high. We find that intermediate switching rates maximize the rate of adaptation on rugged fitness landscapes. This is because an intermediate switching rate can maintain within the same individuals both a mutator phenotype and pre-existing mutations, a combination that facilitates the crossing of fitness valleys. Further, intermediate switching rates allow the population to quickly revert to a low mutation rate after adaptation is achieved, avoiding the accumulation of deleterious mutations linked to mutator alleles. Our results rationalize recently observed noise in the expression of proteins that affect the mutation rate and suggest that non-genetic inheritance of this phenotype may facilitate evolutionary adaptive processes.

## Introduction

### Variation in mutation rates

Mutation rates vary widely between species, ranging from 10^−11^ per bp per generation in *Paramecium tetraurelia* [1] to 10^−5^ per bp per generation in single-strand RNA viruses [2]. They also vary between and within populations [3–5]. This is sometimes due to an increased mutation rate induced by mutator alleles, such *MutS, MutL* [6], *mutT* [7], and *dnaQ* [4] in *Escherichia coli*, which increase the mutation rate by up to 100-1,000-fold.

Interestingly, high variation in mutation rate has also been observed between environmental conditions. This variation can be regulated by environmental or cellular conditions [8] in a phenomenon called s*tress-induced mutagenesis*, which is the increase in mutation rate as a response to stress or maladaptation [9–12]. Stress-induced mutagenesis has been extensively studied in *E. coli* [13], and has also been described in other organisms, including yeast [14], flies [15], and human cancer cells [11,16]. It has been suggested that stress-induced mutagenesis facilitates complex adaptive evolution, increases the population mean fitness at the mutation-selection balance, and can evolve in both constant and changing environments, even in the presence of rare horizontal gene transfer [16–19].

### Evolution of the mutation rate

Mutators—individuals with an above-average mutation rate— often arise spontaneously in evolving populations [7]. Mutators may facilitate adaption yet they become burdened with a higher mutational load compared to their non-mutator counterparts [20,21]. Hence, we expect a reduction of the mutation rate after the population is well-adapted to its environmental conditions, assuming high heritability of the mutator phenotype [7,22,23], such that the mutation rate will evolve towards some physiologically attainable minimum. However, the evolution of the mutation rate may also be affected by genetic drift [24,25], biological and physical constraints [26–28], the cost of replication fidelity [27,29], and second-order selection due to hitchhiking with beneficial mutations [7,30–34]. Additionally, the shape of the fitness landscape might constrain the evolution of mutators [35]. Mutators may also be favoured in situations where beneficial mutations are frequent, such as frequently changing environments that require constant generation of adaptations [36–39]. Frequent generation of mutator alleles could also lead to a substantial equilibrium frequency of mutator alleles, even under strong selection [40]. Indeed, populations evolving from a single ancestor have been found to exhibit variation in the mutation rate, suggesting that the generation of mutator alleles of various intensities may be frequent [4,41]. Experimental validation for the hypothesis that weak mutator alleles could hitch-hike with beneficial mutations in a substantial proportion of cases [30] comes from the *Long-Term Evolutionary Experiment* [33], in which 3 of 12 *E. coli* lineages evolved a 10-100-fold increase in mutation rate after 10,000 generations due to fixation of mutator alleles [33]. However, the mutation rate was later reduced by anti-mutator alleles, perhaps because the opportunities for adaptive evolution diminished [7].

### Non-genetic inheritance of a mutator phenotype

The mutator phenotype is commonly thought to be stable over generations due to genetic inheritance, the result of, for example, frame-shift, deletion, or short-tandem repeat mutations in DNA mismatch repair genes [6,7,42] or DNA polymerase [4]. In contrast, the possibility of non-genetic inheritance of the mutation rate has received little attention. Ninio [43] has suggested that *transient mutators* could arise, for example, as a result of a translation error in DNA repair proteins that could lead to a very strong mutator phenotype. This mutator phenotype could be inherited for a few generations via cytoplasmic inheritance, despite not being encoded in the genotype. In this scenario, individuals can increase their mutation rate for a few generations, thus generating beneficial mutations without a long-term commitment to an elevated mutation rate and the accumulation of deleterious mutations it entails. Rosenberg et al. [44] have also suggested that most adaptive mutations could be due to transient mutators. A more extreme case in which that mutation rate is not inherited, but rather randomly determined, was recently studied by Bonhoeffer et al. [45]. They found that fluctuations in the mutation rate increase the population mean fitness by producing a subpopulation with below-average mutation rate when the population is well-adapted, and a subpopulation with above-average mutation rate when there is potential for further adaptation. In a subsequent paper, it was suggested that this variation could increase population evolvability [46].

There are several lines of empirical evidence for non-genetic inheritance of the mutation rate. Expression noise in genes involved in mutation rate regulation may cause it to fluctuate stochastically. Such noise has been demonstrated for many gene promoters [47] and has been shown to scale with the abundance of the corresponding protein [48]. Translation errors occur more often in some genes involved in DNA repair and replication [49], potentially realizing the scenario proposed by Ninio [43]. Moreover, a mechanism for epigenetic inheritance of the mutation rate in *E. coli* has been described by Uphoff *et al*. [50]: Ada is a DNA repair protein induced under high pH. Each cell carries, on average, a single Ada molecule. Therefore, during cell division a substantial proportion of daughter cells inherit zero Ada molecules, and thus experience an elevated mutation rate. Hence, stochastic fluctuations in Ada copy number can have effects on the mutation rate. Furthermore, *Ada* is positively auto regulated through a mechanism which can intensify the difference in mutation rate between cells that did or did not inherit Ada molecules.

### The inheritance mode of the mutation rate

As a special case of the Adaptation Spectrum of inheritance [51] we suggest a spectrum of mutation arte inheritance: at one end, mutator alleles that arise as rare mutations in genes involved in DNA repair or replication are inherited with very high fidelity, with little to no effect of stochastic factors; on the other end of the spectrum, frequent and stochastic fluctuations in concentrations of proteins that affect the mutation rate may be of such magnitude that there is effectively no correlation between parent and offspring mutation rate; and in the middle of the spectrum, is the range that we define here to be the “epigenetic modes of inheritance”, e.g. of the type that can be attained through cytoplasmic inheritance of proteins or transgenerational epigenetic inheritance of fluctuations in protein concentrations. Such phenomena may produce partially heritable mutator phenotype that is transmitted across cellular generation at an intermediate fidelity. Another potential mechanism is aneuploidy, which arises at rates much higher than genetic mutations, and may produce a significant mutator phenotype [52].

### Crossing fitness valleys

We focus here on an open problem in adaptive evolution, initially presented by Sewall Wright in 1931 [53,54]: if different alleles are separately deleterious but jointly advantageous, how can a population evolve from one co-adapted allele combination to a fitter one, crossing a less fit “valley”? Wright’s solution to this problem, the *shifting balance theory of evolution*, utilizes genetic drift in small populations with limited migration, and is valid [55], but probably limited to specific scenarios [56,57]. Other solutions were proposed, including increased phenotypic variance after population bottlenecks [58], environmental heterogeneity [59], stochastic tunnelling in large asexual populations [60], and stress-induced mutagenesis [16].

### Overview

Here, we investigate adaptation on rugged fitness landscapes with different modes of inheritance of the population variable mutation rate. We have developed a Wright-Fisher model with explicit inheritance of two mutation rate phenotypes: non-mutator and mutator. The phenotype switching rate determines the probability that an offspring will switch from its parents mutation rate phenotype rather than inherit it from its parent. Our model allows to examine rate of adaptation under various switching rates. We find that intermediate switching rates, which correspond to epigenetic inheritance of the mutation rate, lead to the highest rates of adaptation on theoretical and empirical rugged fitness landscapes. We suggest this advantage of intermediate switching rate is conferred upon it since under this regime the mutators that already possess needed genetic mutations towards the ultimate solution is maximized. This combination of mutators with mutations accelerates the rate of fitness-valley crossing on rugged fitness landscapes.

## Models

We first describe our general model, followed by three specific models, each focusing on a different fitness landscape.

### General evolutionary model

We consider an asexual haploid population with non-overlapping generations. We model the effects of mutation, phenotypic switching, natural selection, and genetic drift using a Wright-Fisher model [61]. Individuals are fully characterized by their mutation rate phenotype, either non-mutator (*m*) or mutator (*M*), and their genotype, which determines their fitness. The genotype is specified differently in each fitness landscape (see below). The combination of these two characteristics is termed *pheno-genotype* [62] to emphasize that it is defined through a combination of a non-genetic and a genetic component, namely the mutator phenotype and mutations. The frequency of individuals with mutator phenotype *z* and genotype *g* is denoted by *f*_*zg*_. In all cases we assume no recombination and, therefore, complete linkage.

### Mutation

The pheno-genotype frequencies after mutation are given by

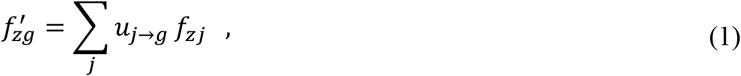

where *j* is an index over all possible genotypes, *z* is the mutation rate phenotype, either *m* or *M* for non-mutator and mutator, respectively, and *u*_*j→****g***_ is the mutation transition probability from genotype *j* to genotype *g*. The specific transition probabilities *u*_*j→****g***_ in each fitness landscape are given below.

### Phenotype switching

In every generation an individual may switch its mutation rate phenotype with probability *γ*. For simplicity, we assume this *switching rate γ* is symmetric from non-mutator *m* to mutator *M* and vice versa (see **Figure 1A**). Thus, the pheno-genotype frequencies after phenotype switching are given by

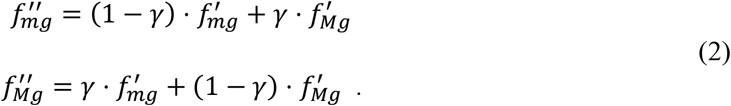

Later, we also extend our model to include a transition pheno-genotype with mutation rate that is intermediate between the non-mutator mutation rate and the mutator mutation rate (see **Figure 1A**).

**Figure 1.**
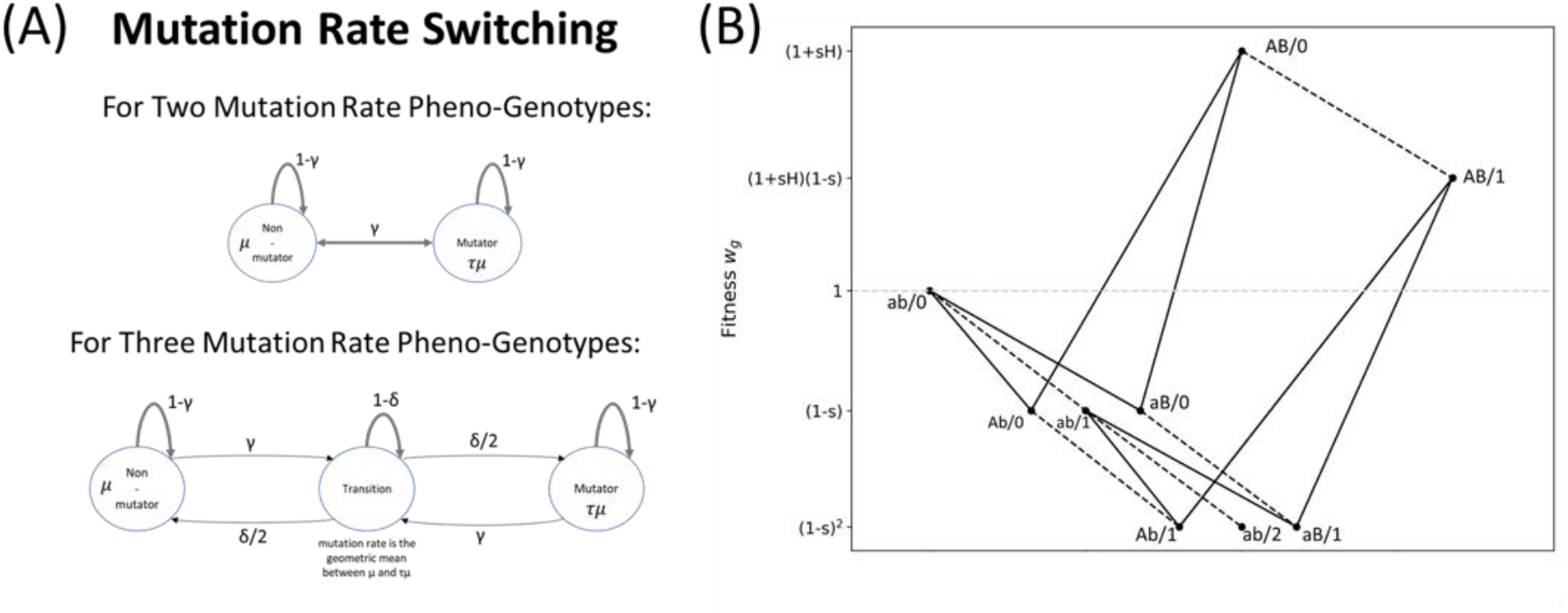
Model illustration. **(A)** With two phenotypes, a cell transitions from a non-mutator (mutation rate *μ*) to a mutator (mutation rate *τμ>μ*) phenotype with probability *γ*, the phenotype switching rate. With three phenotypes, there is a transition phenotype which non-mutator and mutators enter with probability *γ* and leave to either phenotype with probability *δ/2*. **(B)** The two-peak fitness landscape. Fitness (after an environmental change) is shown on the y-axis. Solid and dashed lines denote mutations in major and background loci, respectively. See Table 2 for summary of model parameters. Figure adapted from [16].

### Selection

In accordance with previous work on the evolution of modifiers, we assume that the mutator phenotype is neutral [23,40,63], that is, it does not affect fitness (also, in contrast to most previous work, we do not assume it is an allele, i.e. we do not assume genetic inheritance). Hence, we can focus on the fitness *w*_*g*_ of genotype *g*, rather than the fitness of the pheno-genotypes. The specific fitness values are determined by the fitness landscape, see below. The pheno-genotype frequencies after selection is

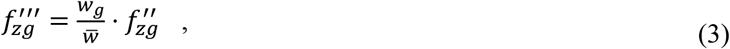

where 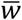 is the population mean fitness,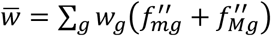.

### Genetic drift

We model the effect of random genetic drift by drawing the number of individuals of each pheno-genotype from a multinomial distribution parameterized by the population size *N* and the pheno-genotype frequencies after mutation, phenotype switching, and selection,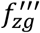.

### Numerical analysis

The effects of random drift can sometimes be neglected (e.g., very large population size). In these cases, pheno-genotype frequencies are only affected by mutation, phenotype switching, and selection. We refer to this version of the model as the *deterministic model* and solve it numerically by iterating eqs. (1-3) until the population reaches an equilibrium (i.e., the population mean fitness does not change from one generation to the next). The equilibrium frequencies can also be obtained by numerically solving for the eigenvalues of the mutation-selection transition matrix, and normalizing the eigenvector corresponding to the leading eigenvalue. We observed that the values obtained by solving the eigenvalue problem corresponded closely to the values obtained from a numerical iteration of the model. The full model with drift is referred to as the *stochastic model*. It is simulated by iteration of eqs. (1-3) and random sampling at each generation using a multinomial distribution. We estimate the appearance or adaptation rate from 500 simulations of the full model using an augmented MCMC scheme, see *Appendix A*.

### Adaptive evolution

We first consider populations at a mutation-selection balance around a local fitness peak. That is, at the mutation-selection balance the population is well-adapted to its environment and the frequencies of the pheno-genotypes do not change from generation to generation 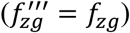 due to a balance between mutation and phenotype switching, which generate genetic and phenotypic variation, and selection and/or drift, which eliminate variation.

In our analytic approximations we assume that selective sweeps [64–67] occur sequentially rather than concurrently, that is, one adaptive genotype goes to fixation before another appears and there is no clonal interference. This assumption is supported by some empirical studies, and corresponds to a regime where selection is strong and mutation is weak (SSWM) [68]. In this SSWM regime, mutations are rare and the advantage they confer is large, i.e., *s>τU* where *s* is the selection coefficient and *τU* is the genomic deleterious mutation rate in mutators. Hence, frequencies of pheno-genotypes with more than one mutation can be neglected at the mutation-selection balance [69]. Based on estimates from the literature (Table 2), 0.001<*s*<0.03, 0.0004<*U*<0.003, and 10<*τ*<1,000. Thus, for the lower range of *s* and higher range of *τU*, our assumption may be invalid.

We consider two types of adaptive evolution dynamics: the crossing of a single fitness valley, and the crossing of multiple fitness valleys until reaching the global fitness peak. In the first case, we approximate the rate of appearance and rate of fixation of an adaptive genotype. From these probabilities, we estimate the adaptation rate of the population and compare it to results of simulations of both the deterministic and the stochastic models. In the case of adaptation to the global fitness peak, we simulate the stochastic model and allow the population to evolve for 50,000 generations, during which we record the population mean fitness and the frequency of non-mutators and mutators in the population.

In the following, we describe three fitness landscapes. In each landscape we define how genotypes are represented and how the pheno-genotype frequencies are affected by mutation (eq. 1) and selection (eq. 3).

### Two-peak fitness landscape

#### Genotype

Here the genotype contains two major loci, in which beneficial mutations may occur, and a large number of background loci, in which only deleterious mutations may occur. Thus, the genotype is denoted by the alleles at the two bi-allelic major loci and by the number of deleterious mutations accumulated in the background loci. For example, the wild type genotype is *ab/0*: it carries alleles *a* and *b* at the major loci and zero mutant alleles in the background loci, while a genotype *aB/2* has the non-wild type allele in the second major locus, B, and two mutations in the background loci.

#### Selection

The fitness of the wild type genotype *w*_*ab/0*_ is set to 1. We assume that all mutant alleles in the background loci are deleterious with a selection coefficient *s*, such that the multiplicative fitness effect of *k* mutant alleles is *(1-s)*^*k*^. Similarly, single mutations at the major loci are deleterious with the same selection coefficient *s*. Initially, the double mutant *AB* is maladaptive with fitness effect *(1-s)*^*2*^. However, after an environmental change occurs, *AB* is adaptive, with fitness effect *(1+sH)*, where *H* is the adaptation coefficient. Hence, after the environmental change there is *sign epistasis* between the two major loci, and the population must cross a *fitness valley* to adapt [54,70]. **Table 1** and **Figure 1B** summarize the fitness values of the different genotypes.

**Table 1.**
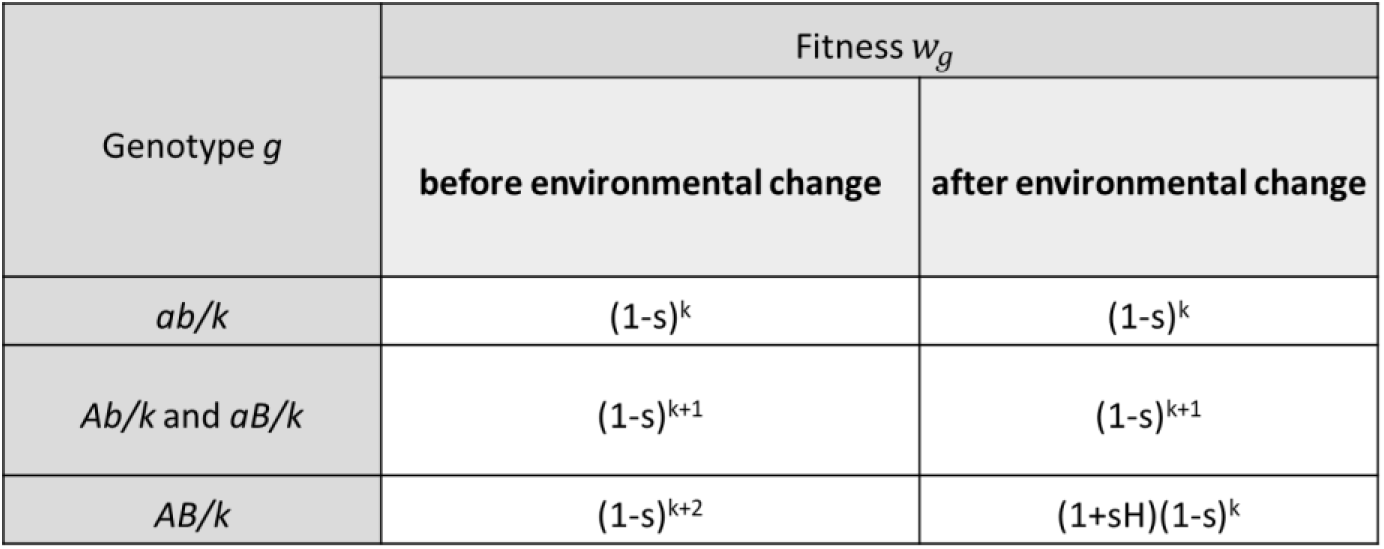
Fitness values in the two-peak fitness landscape. *A, B* and *a, b* are the mutant and wild type alleles of the major loci, respectively; *k* is the number of mutant alleles in the background loci; *s* is the selection coefficient against mutant alleles; *H* is the adaptation coefficient. See Table 2 for summary of model parameters.

**Table 2.** Summary of model parameters. Adapted from [16].

| Symbol | Name | Estimate | References |
| --- | --- | --- | --- |
| $N$ | Population size | $10^5$-$10^8$ | [71] |
| $U$ | Genome-wide mutation rate | 0.0004-0.003 | [72] |
| $n$ | Number of loci | 4,000-5,000 | [73] |
| $\mu$ | Per-locus mutation rate | $U/n$ | |
| $\tau$ | Fold-Increase in mutator mutation rate | 10-1,000 | [32,40,74] |
| $s$ | Selection coefficient against deleterious mutations | 0.001–0.03 | [75] |
| $H$ | Adaptation coefficient | 1–10 | [75] |
| $\gamma$ | Mutator phenotype switching rate | $10^{-6}$-0.5 | [40,43] |
| $\delta$ | Switching rate from transition mutation rate phenotype | 0-1 | |

#### Mutation

The number of mutations per generation that occur at the background loci is Poisson distributed with expected value *U* or *τU* in individuals with non-mutator or mutator phenotype, respectively. Therefore, a genotype with *k* deleterious mutations will mutate to have *k+l* deleterious (*l* is assumed to be positive due to neglecting back mutations) mutations with probability *e*^−*U*^*U*^*l*^/*l*! or *e*^−*τU*^(*τU*)^*l*^/*l*! in individuals with non-mutator or mutator phenotype, respectively. Mutations occur at the major loci with probability *μ = U/n* or *μ = τU/n*, according to the phenotype, where *n* is the total number of loci in the genome and *μ* is the per-locus mutation rate. Therefore, genotype *ab* mutates to *Ab* or *aB* with probability *μ(1-μ)* and to *AB* with probability *μ*^*2*^, and genotypes *Ab* and *aB* mutate to genotype *AB* with probability *μ*. We neglect the effects of back mutations. See *Appendix B* for a formal description of the mutation transition probabilities.

#### Adaptation

The population is initially at a mutation-selection balance around the wild type genotype *ab*/0, which is the fittest genotype before the environmental change. We assume the population size *N* to be large enough so that single mutants (*Ab* and *aB*) are present even in a population of non-mutators, but small enough so that double mutants (*AB*) are absent even in a population of mutators. Thus, the adaptive genotype must appear due to a de-novo mutation rather than from standing variation. Taken together, we assume 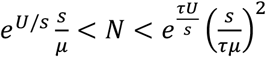 (*Appendix C*). **Table 2** summarizes the model parameters with biologically relevant values.

### Gradual transition between mutator and non-mutator phenotypes

While we mostly consider a model with two mutation rate phenotypes: a mutator and a non-mutator we extended the model to include also an intermediate value of mutation rate. In biological systems, the transition between a mutator and a non-mutator phenotype may be gradual rather than abrupt, and it may potentially occur over several generations (i.e., cell divisions). We therefore extend our model to consider an intermediate mutation rate that is set as the geometric average of the non-mutator and the mutator mutation rates. We call this additional mutation rate pheno-genotype *t*, with *f*_*tg*_ the frequency of an individual with genotype *g* and the transient mutation rate phenotype *t*. We introduce a parameter *δ* that modulates the transition between mutator and non-mutator; the transient mutation rate phenotype (*t*) is inherited with probability *1-δ*, and switches to the mutator or non-mutator phenotype with probability *δ/2* (see **Figure 1A**). Thus, the pheno-genotype frequencies after phenotype switching are given by

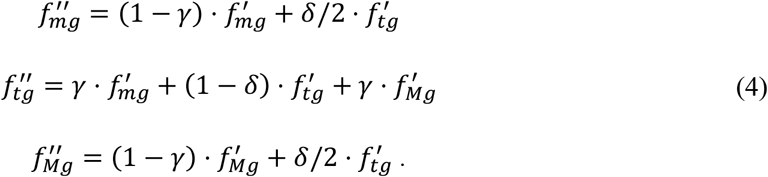

### NK fitness landscape

#### Genotype

The NK landscape is commonly used in the study of epistatic interactions [76]. It has two parameters: *n* for the number of bi-allelic loci in the genotype and *k* for the number of loci each locus interacts with. The main advantage of the NK landscape is its biological interpretation and its tunable ruggedness [77]. Here, the genotype consists of *n=6* bi-allelic loci, which results in 2^6^=64 genotypes.

#### Selection

To construct an NK landscape, we first generate all possible *k*-bit strings (bit strings of length *k*) and assign to each of them a random fitness value between 0 and 1, sampled from a continuous uniform distribution. Genotype *g* is a *n*-bit string, and its fitness *w*_*g*_ is the sum of the fitness effects of the *k-*bit strings that *g* contains. A locus thus influences the genotype fitness according to the number of *k-*bit strings that contain it, which increases with *k*. Hence, the ruggedness of the NK landscape increases with *k* [78]. We consider three NK landscapes with low, intermediate, and high ruggedness corresponding respectively to *k* = 1, *k* = 3, and *k* = 5. See **Figs. A11** and **A12** and **Table A2** for properties of the constructed landscapes.

#### Mutation

A mutation switches the identity of a random bi-allelic locus from one allele to the other. The mutation transition probability *u*_*j→g*_ between two genotypes *j* and *g* that differ in *d* loci is *f(d)*, where the *f* is the probability mass function of a binomial distribution *Bin(n, μ)* for non-mutators or *Bin(n, τμ)* for mutators, *μ* is the per-locus mutation rate, and *τ* is the mutation rate fold-increase in mutators.

#### Adaptation

The initial population is at a mutation-selection balance around a genotype that is a local fitness maximum (see *Appendix D*). In the set of NK landscapes that we generated for the least rugged landscape, *k* = 1, there exists only one local maximum other than the global maximum. This local maximum genotype was at a distance of *d*=4 single-locus mutations from the global maximum genotype. For landscapes of intermediate and high ruggedness with *k=3* and *k=5*, respectively, we also set the initial genotype to be a local maximum at a distance of *d=4* from the global maximum.

### Empirical fitness landscape

We examine the insights gained on two-peak and NK landscapes with an empirical fitness landscape [76]: measured with mutants in 8 genes of *Aspergillus niger* [78]. We chose this landscape because: (i) it is complete with fitness measurements of all possible 256 genotype combinations, so we can avoid the interpolation of fitness values of missing genotypes; and (ii) fitness was measured as growth rate relative to the wild type, as opposed to other studies that quantify some proxy phenotype for fitness such as fluorescence or DNA binding [79–81].

#### Genotype

The genotype consists of eight bi-allelic loci, each with either the wild type or the mutant allele. de Visser *et al*. [78] engineered 256 genotypes to bear all possible combinations of wild type/mutant alleles in these eight loci. The loci are: *fwnA1* (fawn-colored conidiospores); five auxotrophic markers, *argH12* (arginine deficiency), *pyrA5* (pyrimidine deficiency), *leuA1* (leucine deficiency), *pheA1* (phenyl-alanine deficiency), and *lysD25* (lysine deficiency); and two resistances, *oliC2* (oligomycin resistance) and *crnB12* (chlorate resistance).

#### Selection

de Visser *et al*. [78] estimated fitness by measuring the growth rate of each strain relative to the growth rate of the wild type strain (i.e., with eight wild type alleles). Out of 256 genotypes, 70 genotypes were lethal (fitness is zero). The landscape is rugged: it contains 15 local maxima (including the global maximum). See **Figs. A9** and **A10** for the distribution of fitness values.

#### Mutation

As in the NK landscape, a mutation switches a random bi-allelic locus from one allele to the other. The mutation transition probability *u*_*j→g*_ between two genotypes *j* and *g* that differ in *d* loci is again *f(d)*, where the *f* is the probability mass function of a binomial distribution *Bin(n, μ)* for non-mutators or *Bin(n, τμ)* for mutators, *μ* is the per-locus mutation rate, *τ* is the mutation rate fold-increase in mutators, and *n* = 8 is the number of loci.

#### Initialization

The initial population is at a mutation-selection balance around a local maximum genotype (see *Appendix D*) chosen to be *d=4* mutations away from the global maximum.

## Results

### Adaptation on two-peak fitness landscapes

To analyze the adaptation rate, we focus on four pheno-genotypes (their frequencies appear in parentheses): non-mutators with wild type genotype *ab/0* (*m*_*0*_); non-mutators with a single major locus mutant allele, *aB/0* or *Ab/0* (*m*_*1*_); mutators with wild type genotypes (*M*_*0*_) and mutators with a single major locus mutant allele (*M*_*1*_). Note that a single mutant allele is always deleterious in the two-peak landscape. We approximate the mutation-selection balance (MSB) frequencies of these pheno-genotypes in *Appendix E* and compare these approximations to numerical results of the deterministic and stochastic models. We find a good agreement between all three results.

Starting from the MSB, we assume an environmental change provides an adaptive opportunity. In particular, under the new environmental conditions, the double mutant *AB* has the highest fitness. The rate of adaptation is calculated from the probability of appearance and the probability of fixation of this fittest double mutant. These probabilities are derived from the frequencies *m*_*0*_, *m*_*1*_, *M*_*0*_, and *M*_*1*_. We neglect the frequency of a double mutant at the MSB (i.e., we disregard terms of order *μ*^*2*^) due to our assumption on the population size and the mutation rate (see above). Hence, the probability *q* of appearance of a double mutant without background mutant alleles, *AB/0*, as a result of a mutation in a single mutant (either *aB/0* or *Ab/0*) in a population that currently does not have any double mutants is

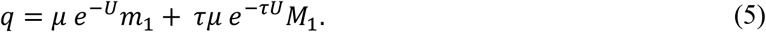

Here, *μm*_*1*_ and *τμM*_*1*_ are the probabilities that a major locus mutation occurs in a single mutant genotype with either the non-mutator or the mutator phenotype, respectively, and *e*^*-U*^ and *e*^*-τU*^ are the probabilities that no background loci mutations occur, again with either the non-mutator or mutator phenotype. Following Eshel [82], the fixation probability *π*of a double mutant once it appears in a large population only depends on the selection coefficient *s* and the adaptation coefficient *H*, 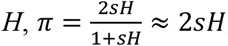, where the latter approximation applies when *sH* is much smaller than 1 (**Figure A5**).

The probability that in the next generation no double mutant appears and fixes in a population of size *N* is (1 − *qπ*)^*N*^. Assuming the number of generations until the appearance of a double mutant that goes to fixation is geometrically distributed, the *adaptation rate v* is defined as the probability that a single double mutant appears and escapes extinction by genetic drift,

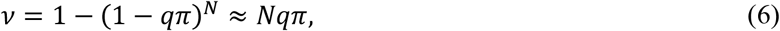

where the approximation holds when *Nqπ* is small. Note that the switching rate *γ* affects the adaptation rate only indirectly due to its role in the MSB frequencies *m*_*0*_, *m*_*1*_, *M*_*0*_, and *M*_*1*_.

We compare results from (**Figure 2**): (i) simulations of the stochastic model (i.e., with background loci and genetic drift); (ii) the approximation in Eq. 6 using MSB frequencies from our analytic approximation (*Appendix E*); and (iii) the approximation in Eq. 6 using numerically determined MSB frequencies (without drift, Eqs. 1-3). We find good agreement between the two approximations, (ii) and (iii) (*R*^*2*^=0.96), as well as between (i) and (ii) (*R*^*2*^=0.93) and (i) and (iii) (*R*^*2*^=0.96). This is despite our analytic approximation assuming only four genotypes and no drift, which neglects background effects such as genetic hitchhiking and clonal interference (the stochastic model includes drift as well as background effects). We plotted the appearance rate (Eq. 5) vs. the switching rate γ in **Figure 2A**, and the adaptation rate (Eq. 6) vs. the switching rate γ in **Figure A6**.

**Figure 2.**
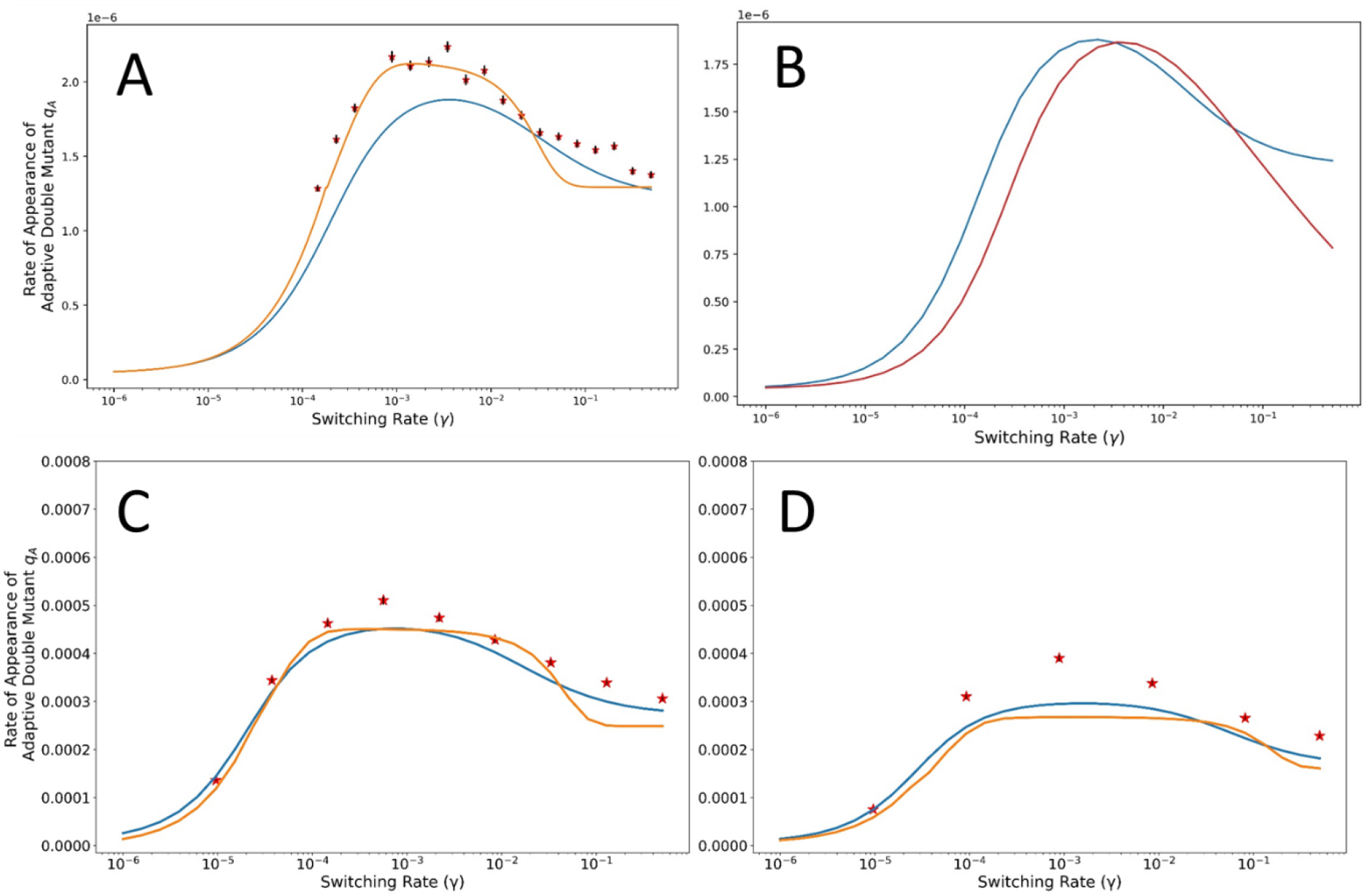
Intermediate switching rates maximize adaptation rates. Stars show the appearance rate estimated from simulations (the stochastic model with genetic drift). Lines show the approximation (Eq. 6) using MSB frequencies from an analytic approximation (orange; Appendix E, Eqs. A3-A16) or numerical analysis (blue; Eqs. 1-3 without drift). **(A)** Two-peak landscape with two mutation rate phenotypes, mutation rate, *μ=4·10*^*-5*^; mutation rate increase, *τ=10*; selection coefficient, *s=0.03*; adaptation coefficient, H=5, population size *N=10*^*7*^. **(B)** Two-peak landscape with two (blue) or three (red) mutation rate phenotypes (see **Figure** 1A), *δ=0.5*. **(C)** NK landscape, *k=3*; *s=0.03* (average effect of deleterious mutations on the fitness, see *Appendix H*); *N=10*^*6*^. **(D)** Empirical landscape, *μ=10*^*-6*^, *s=0.15* (average effect of deleterious mutations on the fitness, see *Appendix H), N=10*^*5*^. Coefficients of determination, *R*^*2*^, for every combination of orange, blue lines and stars are shown in Table A3 and are all greater than 0.9. Error bars represent 90% CI, see *Appendix A*.

Importantly, the highest appearance and adaptation rate is reached with intermediate switching rates (**Figure 2**). This result applies for a wide range of mutation rates: we find that intermediate switching rates in the range of 0.001<*γ*<0.01 maximize the rate of adaptation for a wide range of parameters (**Figure 3**). To explain the advantage of intermediate switching rates, we note that the frequency of mutator mutants *M*_*1*_ has a larger effect on the appearance probability *q*_*A*_ compared to the frequency of non-mutator mutants (Eq. 5, because *U<1*). Thus, to determine the role of the switching rate *γ* on the adaptation rate, we must determine how it affects *M*_*1*_. Indeed, we find that *M*_*1*_ is maximized when the switching rate is intermediate (**Figure A4D**), namely (*Appendix F*)

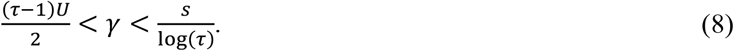

This is because *M*_*1*_ is approximately the ratio between a logistic function with rate *(τ-1)U/2*, describing the proportion of mutants at MSB (*m*_*1*_*+M*_*1*_), and a logistic function with rate *s/log(τ)*, describing the decrease in the co-occurrence between the mutator phenotype and the mutant genotype (*m*_*1*_*/m*_*0*_; *Appendix E*). A heat-map of upper bound and lower bound on *γ* versus the relevant parameters is shown in **Figure A8**.

**Figure 3.**
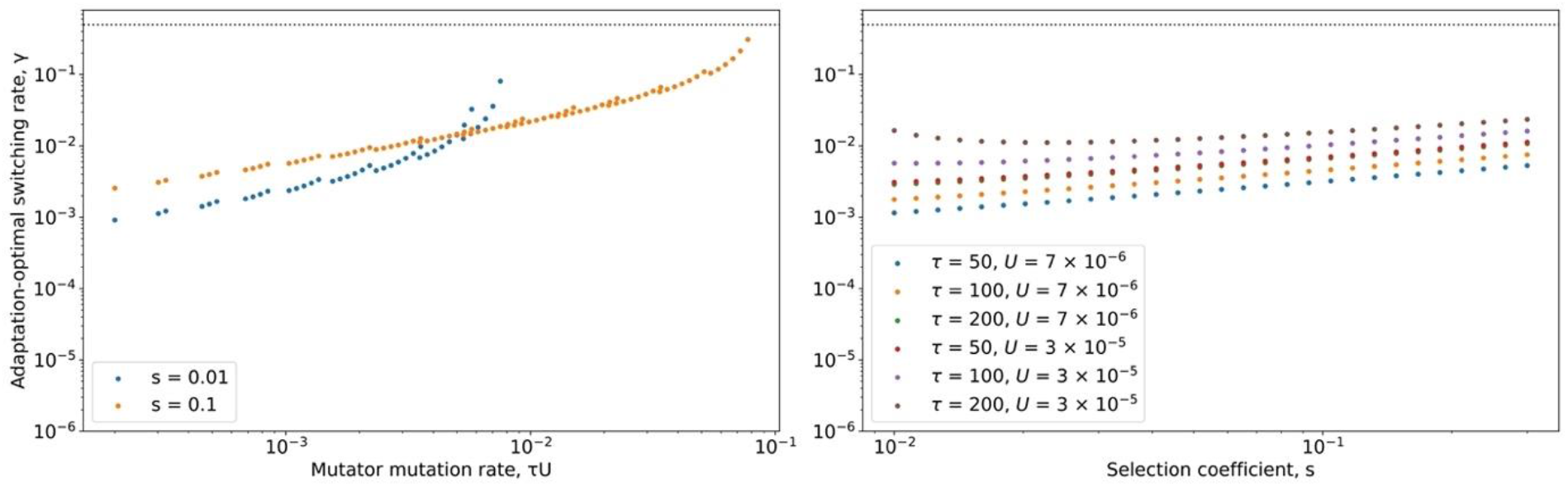
Adaptation-optimal switching rates. The adaptation-optimal switching rate **(A)** increases with the mutator mutation rate, *τU*, until reaching *γ*=0.5 when *τU=s*; and **(B)** is marginally dependent on the selection coefficient *s*. The optimal switching rate *γ* maximizes the double mutant appearance rate (Eq. 5, ***q*** = ***μ e***^−***U***^***m***_**1**_ **+ *τμ e***^−***τU***^***M***_**1**_) and was found using *scipy.minimize* (5). *m*_*0*_, *m*_*1*_, *M*_*0*_ and *M*_*1*_ computed by numerical iteration of the model on the two-peak fitness landscape. Here, selection coefficient, *s*=0.01.

### Gradual transition between mutator and non-mutator phenotypes

When considering a third, transient phenotype between the non-mutator and mutator phenotypes, we also observe that the adaptation rate is maximised for intermediate switching rates (**Figure 2B**). The curve describing the adaptation rate as a function of the switching rate is shifted to the right compared to the same curve for the two-phenotype case; this is because the transition rate between non-mutator and mutator and vice-versa is effectively lower at *γδ/2<γ*. See *Appendix G* for the equilibrium frequencies of the phenotypes as a function of the switching rate.

### Adaptation on NK and empirical landscapes

#### Crossing a single fitness valley

In the NK and empirical landscapes, the initial genotype is defined as the wild type while all other genotypes are defined as mutants (see *Appendix D*). To reach a mutation-selection balance (MSB) around the wild type, we temporarily define the wild type to be the genotype with the highest (global maximum) fitness (*Appendix D*), and so the frequency of mutants was low compared to that of the wild type. After reaching the MSB, the fitness values of the mutants were reset to their landscape values.

First, we evaluate the rate of crossing a single fitness valley as in the case of a two-peak landscape. The rate of appearance of the double mutant (i.e., a mutant at a higher fitness peak two mutations away from the wild type) was approximated using Eq. 5 with MSB frequencies *m*_*1*_ and *M*_*1*_ determined numerically (from Eqs. 1-3) or with our analytic approximation, as in the two-peak fitness landscape (*Appendix E*, Eqs. A3-A16). Here, *m*_*1*_ and *M*_*1*_ correspond respectively to the frequencies of non-mutators and mutators with genotypes one mutation away from the wild type. Our approximation thus neglects the frequencies of genotypes more than one mutation away from the wild type. **Figure 2** shows a comparison of these approximations with results of simulations of the stochastic model with drift. Here we focus on the appearance rate because the switching rate does not affect the fixation probability, which is the other factor determining the adaptation rate.

We find that the highest rate of double mutant appearance occurs for an intermediate switching rate. Like in the two-peak landscape, we conclude that intermediate switching rates maximize the rate of crossing fitness valleys in multi-dimensional fitness landscapes such as NK landscapes and the empirical landscape. This conclusion is true regardless of the extent of ruggedness in the NK model (see **Figure A16**).

#### Crossing multiple fitness valleys

Next, we sought to confirm that the advantage of intermediate switching rates extends beyond a single valley crossing. Again, we initialized the population at a MSB around a local maximum of the fitness landscape, which we mark as the wild type (*Appendix D*). We then allowed the population to evolve for 50,000 generations. At each generation, we recorded the proportion of simulations (out of 500) in which the genotype with the highest fitness (the global maximum) became dominant in the population. In the NK landscape with *K*=5, we use the proportion of simulations in which a genotype fitter than the wild type fixed in the population, because none of the simulation reached the global maximum, probably due to the high ruggedness of the landscape.

Again, we find that populations with an intermediate switching rate have the highest adaptation rate (**Figure 4**). We again hypothesized that this advantage is due to the co-occurrence of the mutator phenotype and standing genetic variation, which increases the probability to generate double mutants. To test this hypothesis, we artificially removed this co-occurrence: at each generation, each individual was assigned a mutation rate phenotype at random, independent of its parent phenotype and its genotype, such that the frequency of the phenotypes is equal to their frequency at that generation from a simulation of the stochastic model. With an intermediate switching rate *γ* = 3.4·10^−3^, and with *N*=10^6^, *μ*=10^−6^, and either *τ*=10 (for the NK landscape) or *τ*=100 (for the empirical landscape), the mutator frequency at MSB is almost 50%, close to the mutator frequency at MSB for high switching rate, *γ=0.5*. Therefore, after reassigning mutation rate phenotypes at random among all individuals in the population, we expected the population to evolve similarly to a population with a high switching rate and without phenotype reassignment—that is, at a slower rate compared to intermediate switching rate without phenotype reassignment. Indeed, **Figure 4** shows that populations with intermediate switching rates and random phenotype reassignment (dotted red lines) evolve slower than corresponding populations with intermediate switching rate yet with real phenotype assignment, and similarly to populations with high switching rates (solid orange lines). This result supports our suggestion that fast adaptation with intermediate switching rates is due to the co-occurrence of the mutator phenotype and the mutant genotype.

**Figure 4.**
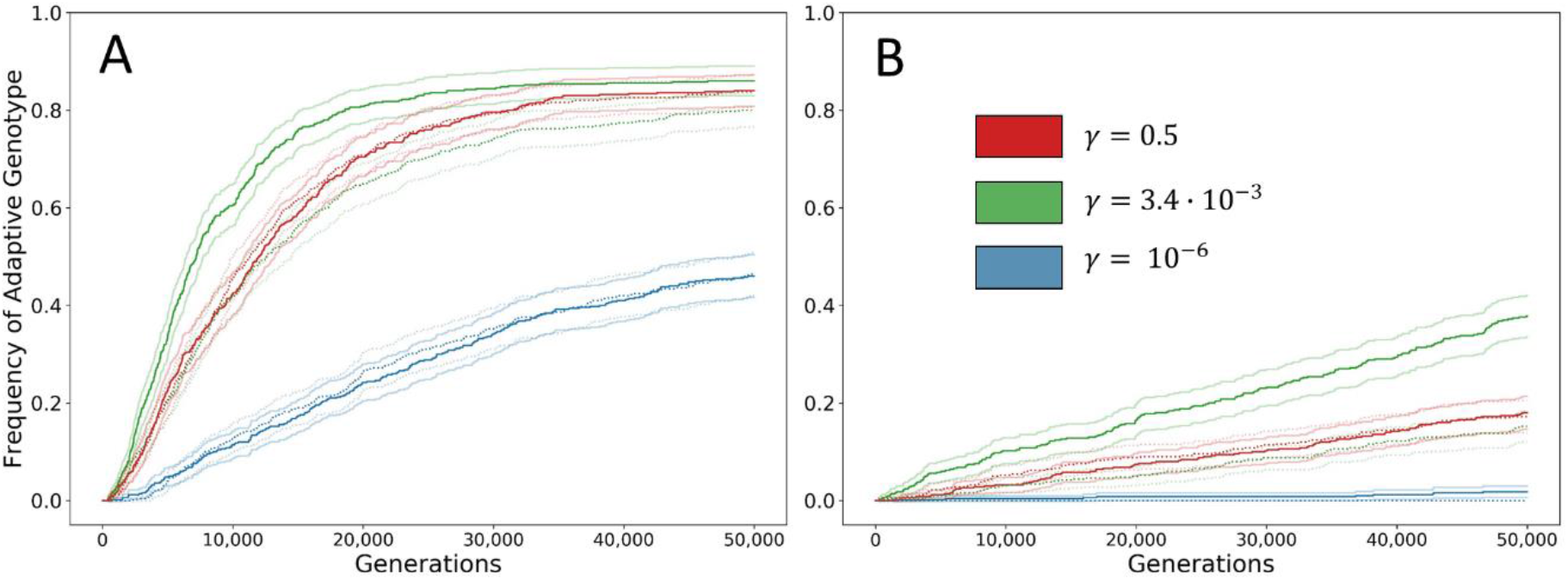
Intermediate switching rates maximize the adaptation rates on NK and empirical fitness landscapes. Solid lines show the frequency of the adaptive genotype (global maximum) from simulations (stochastic model with drift). We observed that intermediate switching rate lead to the highest frequencies of adaptive genotype, and hypothesized that this is due to the co-occurrence of the mutator phenotype with pre-existing genetic variation. To test this hypothesis, we re-ran the simulation while removing the co-occurrence of the mutator and its genotype: at each generation, each individual was re-assigned a random mutator rate phenotype while keeping the mutator frequency equal to the frequency in the original simulation at any given generation (dotted lines). Transparent lines show the corresponding 95% confidence intervals. **(A)** NK landscape with *k=3*; population size, *N*=10^6^; mutation rate, *μ*=10^−6^; mutation rate increase, *τ*=10. **(B)** Empirical fitness landscape; same as A except *τ*=100.

## Discussion

### The advantage of intermediate mutation-rate phenotype switching rates

Through a combination of analytical and computational analysis, we have demonstrated that the adaptation rate on rugged fitness landscapes in populations that consist of mutators and non-mutators is maximized when the rate of inheritance of the mutation-rate phenotype is intermediate, so that is higher than genetic transmission but lower than random determination (i.e., no inheritance). In particular, intermediate rates of mutation-rate phenotype switching, roughly 0.001<*γ*<0.01 (assuming strong selection and weak mutation, i.e., *s>τU*).

With a low switching rate at *γ*=10^−6^, which corresponds to genetic inheritance of the mutation rate, mutators are strongly associated with their mutant alleles, both deleterious and beneficial ones. Therefore, selection efficiently acts against mutators due to the deleterious mutations they generate and accumulate, and their frequency thus declines. On the other side of the spectrum, when the switching rate is high, *γ*=0.5, mutators reach a high frequency; however, they tend not to co-occur in the background of the mutant alleles that they generated. Thus, at such high switching rate the co-occurrence between being a mutator in the background of a mutant that already has the adaptive mutations is quickly broken and the mutators do not accumulate the additional needed mutations. In contrast, intermediate switching rates strike a balance between a high frequency of mutators in the population and a strong co-occurrence of the mutator and the mutant alleles it generates (**Figure 5**). This allows rapid generation of double mutants, which are crucial for adaptation on rugged fitness landscapes. Overall, intermediate switching rates are within the range *(τ-1)U/2<γ<s/log(τ)*.

**Figure 5.**
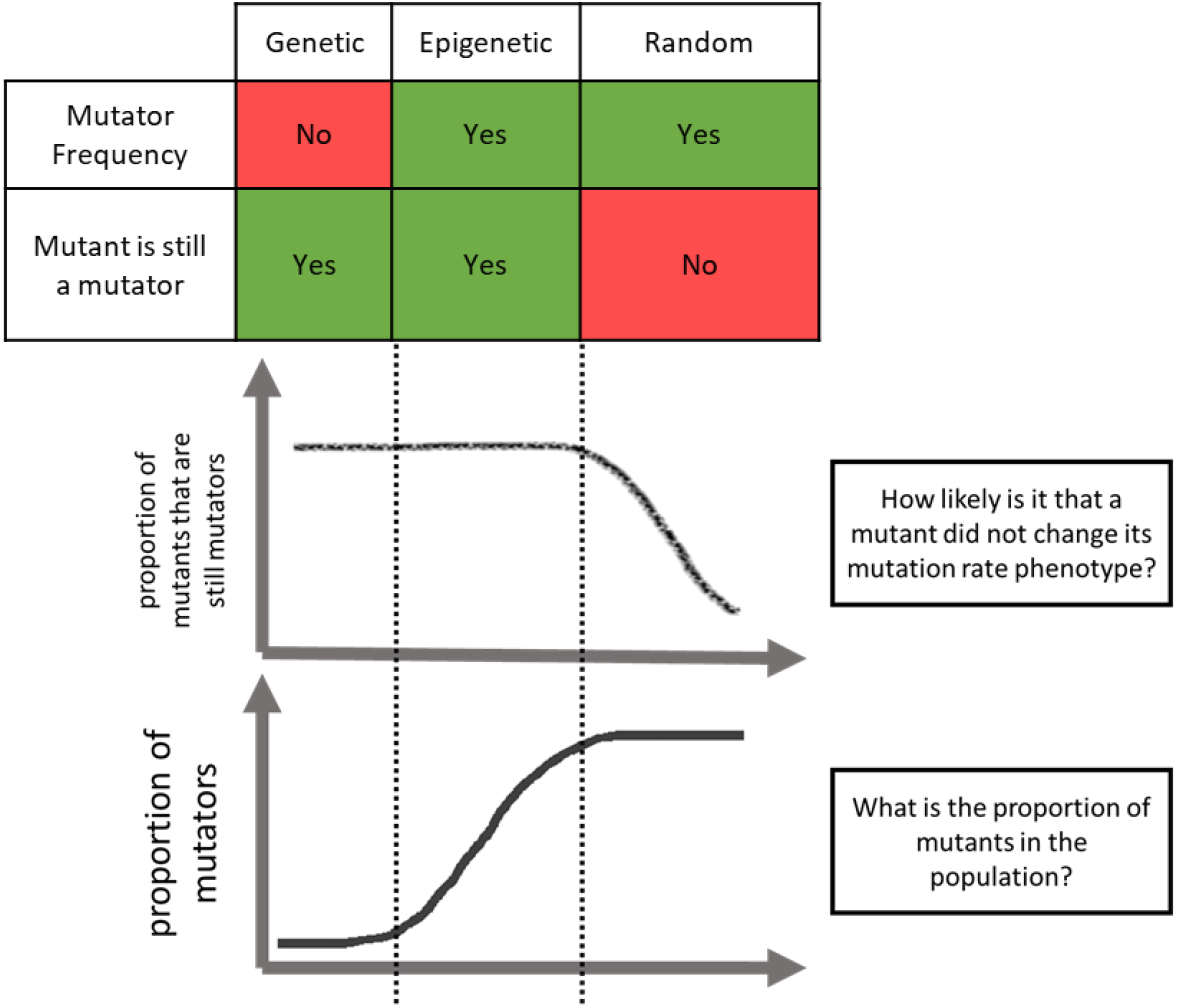
The advantage of intermediate switching rates. Due to the combination of high mutator frequencies at mutation-selection balance and maintenance of the co-occurrence of a mutator phenotype and single mutants, intermediate switching rate maximize adaptation rates on rugged fitness landscapes, where production of double mutants is crucial.

### Mutation rate inheritance and phenotype switching

Switching between a mutator and a non-mutator phenotype is a type of phenotype switching [83]. The increased adaptability of populations with intermediate switching rates, compared to populations with low switching rates, is due to the high rate of mutator generation, which balances the strength of selection against mutators at the mutation-selection balance. Generation of a phenotype that is disadvantageous in the short term but potentially advantageous in the long term is known as *bet-hedging* [83] and has been implicated as driving the evolution of phenotype switching mechanisms [83–86]. The recent demonstration that the DNA repair enzyme Ada in bacteria display noise in expression level, and hence variation in mutation rate in population is an example for realization of such dynamics. Further, the fact that Ada is an auto-regulator, implies that it manifests a non-random inheritance upon cell divisions, and as such it may actually display intermediate switching rate of that type that is shown here to be optimal. The notion of “phenotypic mutators” [43] in which DNA repairs enzymes were suggested to be generated with translation errors followed by inheritance of proteins that harbour such errors, may provide another realization of noisy mutation rate with intermediate heritability. Here, our phenotype of interest is the mutation rate, and the product of the phenotype is the rate of generation of mutations. We find that intermediate switching rates maximize the adaptation rate in rugged fitness landscapes. Likewise, we suggest that intermediate switching rates may be optimal in other systems in which a specific phenotype has a short-term disadvantage, and therefore a high switching rate is required to counter the effect of natural selection. In such cases, if the same phenotype may also be advantageous if it occurs in consecutive generations of the same lineage, and therefore a not too high switching rate is beneficial in maintaining a correlation between the phenotype of parent and offspring.

### Evolution of intermediate switching rates

Although intermediate switching rates maximize the adaptation rate, it is not clear that an adaptation-optimal switching rate can be selected for when competing against lower or higher switching rates. First, the proportion of mutants increases with the switching rate (**Figure A2**). This means that the mutational load of subpopulations with higher switching rates will be higher than the mutational load of subpopulations with lower switching rates, thus selecting for low switching rates. Second, the advantage of a subpopulation with an adaptation-optimal switching rate depends on the mutation supply. The adaptation rate of a subpopulation of size *N*_1_ with a given switching rate *γ*_1_ resulting in an appearance rate *q*_1_ will be higher than the adaptation rate in a subpopulation of size *N*_2_ with a given switching rate *γ*_2_ resulting in an appearance rate *q*_2_ if 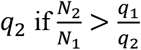 (Eq. 6). Therefore, selection for an allele that induces an intermediate switching rates will depend on its frequency in the population, similar to the case of mutator alleles [87].

For example, in the two-peak landscape with parameter values from **Figure 2**, the highest adaptation rate (with *γ*=3.4·10^−3^) is 36-fold higher than the lowest adaptation rate (with *γ*=10^−6^). Hence, a subpopulation with the adaptation-optimal switching rate might be outcompeted by a subpopulation with a low switching rate if its initial frequency is less than 1/36 ∼ 3%. Note that before the adaptive genotype appears, the frequency of the subpopulation with a high switching rate may decrease due to its higher mutational load, which further decreases its adaptive advantage. A more detailed treatment of the evolution of the adaptation-optimal switching rate is required to conclude whether we could expect to observe selection for intermediate rate of mutation-rate phenotype switching in natural populations.

## Conclusion

Empirical evidence indicates that the mutation rate may be determined both by genetically inherited factors, such as mutator alleles coding for deficient mismatch repair mechanisms [33,88], and by stochastic factors, such as fluctuations in protein concentrations, which may also be epigenetically inherited for a limited number of generations [47,49,50]. We found that the adaptation-optimal switching rates are generally in the epigenetic range and increase with the mutator mutation rate until reaching values that can be interpreted as random determination of the mutator phenotype (**Figure 3**). However, adaptation-optimal switching rates were not found in the range of genetic inheritance. Variance in mutation rates has been previously suggested to increase the population mean fitness both in well-adapted and adapting populations [45]. This variance can be manipulated experimentally [47,89] and hence the interplay between the switching rate and the capacity to adapt could be tested empirically.

## Acknowledgments

YP holds the Ben May Professorial Chair, and is a Kimmel Investigator.

YR acknowledges the Minerva Foundation and the Israel Science Foundation (552/19) for grant support.

We thank Eytan Domany, Ariel Amir, Naama Barkai, and Tal Simon, for discussions and comments.

## Code Availability

Source is available at https://github.com/gabriela3001/phenotypic_mrate.

## Notes

### Competing Interest Statement

The authors have declared no competing interest.

