## Appendix for "Phenotype switching of the mutation rate as a strategy for crossing fitness valleys"

### Appendix A. Augmented MCMC method

We perform 500 simulations of the full model and stop the simulations when 475 of the 500 simulations terminate in order to save computation time. Each simulation terminates either when the adaptive genotype appeared (when focusing on appearance) or when the adaptive genotype exceeded 99% in frequency (when focusing on adaptation). The results – time for appearance (or adaptation) – from 475 simulations are fitted to a geometric distribution. Considering only the first 475 simulations that terminated introduces a Type II Censoring, which complicates fitting a geometric distribution to the data to get the rate of appearance (or adaptation), as using the standard maximum-likelihood estimate:

$$\hat{p} = \frac{n}{\sum_{i=1}^{n} T_i} \quad \text{(A1)}$$

where $T_i$ is the number of generations until appearance or adaptation in simulation $i$, $1 \leq i \leq n$.

We thus apply an augmented MCMC Metropolis-Hasting scheme. We start with a prior estimate of the parameter $p_k$ from $n$=475 samples using eq. A1. Then at each iteration $k$=1,…,5000:

1) Draw the missing 25 samples from a geometric distribution $Geom(p_{k-1}) + \max_{1 \leq i \leq 475} T_i$.
2) Estimate a candidate $p^*$ from the 475 simulations augmented by the 25 samples drawn in step 1 using eq. A1
3) Compute the likelihood $L(p^*)$ on the 500 samples (including simulation results and augmented samples) using the probability mass function of the geometric distribution.
4) With probability $min\left(1, \frac{L(p^*)}{L(p_{k-1})}\right)$, accept the candidate, $p_k = p^*$. Else, reject it and set $p_k = p_{k-1}$.

After discarding a burn-in period of 2,500 iterations, the remaining $\{\hat{p}_k\}_{k=2500}^{5000}$ are considered samples from the posterior distribution of $p$. The mean of this distribution estimates the true parameter of the geometric distribution. A confidence interval is given by the 5$^{th}$ and 95$^{th}$ percentiles of this distribution.

### Appendix B. Transition probabilities in the two-peak landscape

The mutation transition probability is $u_{j\to g}$, where $j$ and $g$ are genotypes, determines the effect of mutation on the genotype probabilities (Eq. 1). The construction of the mutation transition probability is described in the main text. Here, we provide a formal description. Note that we assume no back mutations occur. In the following we use the mutator phenotype mutation rate $\tau U$; for the non-mutator, we set $\tau=1$. The per-locus mutation rate is $\mu = U/L$. The number of deleterious mutations in the background loci is $k = 0,1,2, \ldots$.

Thus, the probability to transition from genotype $j$ to genotype $g$, $u_{j\to g}$, is described by **Table A1**, where the source genotype $j$ is given in the row, and the target genotype $g$ is given in the column. We do not distinguish between *Ab* and *aB* as we assume they have equal fitness values.

| $\boldsymbol{u_{j \to g}}$ | g = ab/k+l | g = Ab/k+l or aB/k+l | g = AB/k+l |
|---|---|---|---|
| j = ab/k | $(1-\tau\mu)^2 \cdot \frac{e^{\tau U}(\tau U)^l}{l!}$ | $2\tau\mu(1-\tau\mu)\frac{e^{\tau U}(\tau U)^l}{l!}$ | $(\tau\mu)^2 \frac{e^{\tau U}(\tau U)^l}{l!}$ |
| j = Ab/k or aB/k | 0 | $(1-\tau\mu) \cdot \frac{e^{\tau U}(\tau U)^l}{l!}$ | $\tau\mu \cdot \frac{e^{\tau U}(\tau U)^l}{l!}$ |
| j = AB/k | 0 | 0 | $\frac{e^{\tau U}(\tau U)^l}{l!}$ |

**Table A1. Probabilities to transition from genotype *j* to genotype *g*, $\boldsymbol{u_{j \to g}}$.**

**Appendix C. Bounds for population size**

We consider $N$, the population size, to be large enough so that individuals with a single mutation in the major loci are present, but small enough so that individuals with two mutations in the major loci (hence already adapted to the new environment) are absent.

Let us first consider the first condition. The lowest frequency of mutants is achieved when the whole population is non-mutator. Therefore, the upper bound for the first condition is set by considering a case where all individuals have the non-mutator phenotype. According to (1), the frequency of wild type individuals at mutation-selection balance is $e^{-U/s}$. We then multiply by the mutation-selection balance of single mutants at the major loci, $\frac{\mu}{s}$. Thus, we find that the expected number of single mutant individuals without deleterious mutations in the background loci is $N\frac{\mu}{s}e^{-U/s}$. Setting this to be less than one and rearranging, we obtain $\frac{s}{\mu}e^{U/s} < N$.

Now let us consider the second condition. The highest frequency of double mutants would be achieved when the whole population is mutator. Therefore, the lower bound for the second condition is set by considering a case where all individuals have the mutator phenotype. By the same argument as for the first condition, we find $N < \left(\frac{s}{\tau\mu}\right)^2 e^{\frac{\tau U}{s}}$ with $e^{\frac{-\tau U}{s}}$ the proportion of wildtype individuals in an all-mutator population and $\left(\frac{\tau\mu}{s}\right)^2$ the frequency of double mutants in the major loci, both at a mutation selection balance.

Combining these conditions, we have

$$\frac{s}{\mu}e^{U/s} < N < \left(\frac{s}{\tau\mu}\right)^2 e^{\frac{\tau U}{s}}. \tag{A2}$$

**Appendix D. MSB initialization for NK and empirical landscape**

First, denote the initial genotype by $g_0$ and its fitness by $w_0$. The set of all single mutants of the initial genotype is $\{g_{1,i}\}_i$, where $i$ indexes the single mutants. The minimum and maximum of the single mutant fitness values are $w_{1,min}=min_i\{w_{1,i}\}$ and $w_{1,max}=max_i\{w_{1,i}\}$. We assume that $w_0>w_{1,max}$, so that the initial genotype by $g_0$ is a local maximum on the fitness landscape.

To obtain a population at mutation-selection balance around $g_0$, we temporarily modify the fitness landscape. The fitness of the initial genotype $w_0$ , as well as the fitness values of its single mutants, $w_{1,I}$, remain the same, but the fitness of all other genotypes is set to $w_{1,min}$. This modification changes the landscape to a smooth single-peak landscape and makes $g_0$ a global maximum. We then iterate the model equations (eqs. 1-3) until a steady state is reached. Then, the simulation can start by restoring the fitness landscape to its full form; that is, the fitness values are restored to their original values.

**Appendix E. Mutation-selection balance frequencies**

In the two-peak fitness landscape, all single mutations in the wild type are deleterious, including at the major loci. Therefore, we consider four pheno-genotypes: non-mutators without deleterious mutations, with frequency $m_0$; non-mutators with a single deleterious mutation, with frequency $m_1$; mutators without deleterious mutations, with frequency $M_0$; and mutators with a single deleterious mutation, with frequency $M_1$. Because mutations in the major loci are as deleterious as mutations in the background loci, we do not distinguish between a mutant in the major loci and a mutant in the background loci. Hence the notations $m_0$, $m_1$, $M_0$ and $M_1$ instead of *Mab/0, mAb/0, Mab/0*, etc.

The change in pheno-genotype frequencies due to mutation, phenotype switching, and selection are described by the following matrix equation,

$$\bar{w} \cdot f' = f \cdot Q \qquad \text{(A3)}$$

where $f = (m_0, m_1, M_0, M_1)$ is the pheno-genotype frequencies vector, $\bar{w}$ is the population mean fitness, and $Q$ is the transition matrix (see **Table 2** for description of parameters). Here,

$$Q = \tilde{Q} D \qquad \text{(A4)}$$

where $\tilde{Q}$ is a right stochastic matrix, i.e. non-negative matrix with rows that sum to 1, and $D$ is a diagonal matrix with positive diagonal entries *1*, *1-s*, *1*, and *1-s* (the fitness values). Therefore, by the Perron-Frobenius theorem (2) the equilibrium of the system in Eq. A3 is characterized by the leading eigenvalue of $Q$ and its corresponding non-negative eigenvector. However, this eigenvalue is complicated and cannot be easily interpreted. We therefore use a simpler approximation.

There is extensive literature on the frequency of *mutators* (3) and *mutants* (1) at a mutation-selection balance, and the pheno-genotype frequencies sum to 1. These quantities are intuitive and can be written as a system of linearly independent equations (*Eq. A5A* below), which can then be solved to obtain the frequencies $m_0$, $m_1$, $M_0$ and $M_1$ at MSB (i.e. the non-negative eigenvector of $Q$).

Hence, we transform the pheno-genotype frequencies to the following variables: the frequency of mutators at equilibrium $p_M$, the frequency of mutants at equilibrium $p_S$, and the ratio of single mutant non-mutators to wild type non-mutators at equilibrium $p_R$,

$$\begin{cases} p_M = M_0 + M_1 \\ p_S = m_1 + M_1 \\ p_R = \dfrac{m_1}{m_0} \\ m_0 + M_0 + m_1 + M_1 = 1. \end{cases} \qquad \text{(A5A)}$$

Rearranging this system, we find expressions for the pheno-genotype frequencies,

$$\begin{cases} m_0 = \dfrac{1 - p_M}{1 + p_R} \\ m_1 = p_R \dfrac{1 - p_M}{1 + p_R} \\ M_0 = p_M - p_S + p_R \dfrac{1 - p_M}{1 + p_R} \\ M_1 = p_S - p_R \dfrac{1 - p_M}{1 + p_R}. \end{cases} \qquad \text{(A5B)}$$

Next, we approximate $p_M$, $p_S$, and $p_R$ as functions of the switching rate, $\gamma$.

**Frequency of Mutators, $p_M$**

The frequency of mutators at the mutation-selection balance has been previously studied by Desai & Fisher (3), abbreviated as DF2011. They considered a scenario where mutators appear due to mutations in, for example, DNA repair genes, and thus they assumed mutators are generated at a very low rate (switching rate $\gamma \approx U \ll 1$, where U is the genomic mutation rate). They also neglected the possibility of switching back from mutator to non-mutator, because when the switching rate is low, mutators will accumulate deleterious mutations and be eliminated from the population before they have a chance to switch back to non-mutators. Therefore, for low switching rates, we use the approximation from DF2011 (Eq. 16 in (3)) to describe the frequency of mutators at equilibrium $p_M$, which is, when $(\tau - 1)U \ll s$,

$$p_M^- = \frac{\gamma}{(\tau - 1)U}. \tag{A6}$$

As the switching rate increases, switching back from mutator to non-mutator can no longer be neglected. The frequency of mutators $p_M$ can then be approximated by the probability that a mutator did not switch before being eliminated due to selection against the deleterious mutations it generated,

$$p_M^+ = \hat{p}_M - \frac{w_M}{\gamma}, \tag{A7}$$

where $\hat{p}_M$ is the proportion of mutators at mutation-selection balance when $\gamma$ = 0.5, and $w_M$ is the mean fitness of mutators at the MSB, determining the expected number of generations mutators survive before being purged by selection. This approximation assumes geometrically distributed waiting times for elimination and switching. When the switching rate is $\gamma = 0.5$, selection cannot act on the mutator phenotype because the phenotype switches, on average, every other generation. Thus, the equilibrium between the mutator and the non-mutator will correspond to the equilibrium of a symmetric two-state Markov chain, whose stationary distribution is the vector $(0.5, 0.5)$. Therefore, we have $\hat{p}_M = 0.5$.

Denote the mutator mutation load, or the mean number of mutant alleles per mutator individual, as $\delta_M$. We approximate the average disadvantage of the mutants with regards to the wildtype by $\delta_M s$ for $s << 1$. We use here $1 - (1 - s)^{\delta} \sim \delta s$. DF2011 derived an expression for the mutational load of mutators at the MSB (Eq. 52 in (3)) ,

$$\delta_M^* = (\tau - 1)U - (\tau - 1)U \frac{(\tau - 1)U}{s}. \tag{A8}$$

Combining Eqs. A7, and A8, we obtain an approximation for $p_M$ at high switching rates,

$$p_M^+ = 0.5 - \frac{\delta_M^* s}{\gamma} = 0.5 - \frac{\left(\tau U - (\tau - 1)\, U \frac{(\tau - 1)U}{s}\right) s}{\gamma} = \frac{\tau U s - (\tau - 1)^2 U^2}{\gamma}. \tag{A9}$$

When the switching rate $\gamma$ is low, the mutator phenotype will be eliminated through second-order selection before it can switch to non-mutator phenotype. When the switching rate $\gamma$ is high, individuals will switch back and forth between phenotypes before selection can affect their frequencies—when $\gamma$ is high enough the trait is not heritable—and $p_M$ will be approximately 0.5.

The transition point between $p_M^-$ and $p_M^+$ will occur in the second (largest) intersection points of $p_M^-$ (eq. A6) and $p_M^+$ (eq. A9). Solving $p_M^+ = p_M^-$ for $\gamma$ using the quadratic formula, this occurs at

$$\gamma_0 = \frac{0.25\, \left(s(\tau - 1) + \sqrt{-s(\tau - 1)(-16U^2\tau^3 + 48U^2\tau^2 - 48U^2\tau + 16U^2 + 15s\tau + s)}\right)}{s}. \tag{A10}$$

In practice, we find that the transition point

$$\tilde{\gamma}_0 = \frac{U(\tau - 1)}{2} \tag{A11}$$

has a simpler expression and results in a good approximation for $p_M$. **Figure A1** shows a comparison between our approximation and numerical results of the deterministic model (Eqs. 1-3 in the main text).

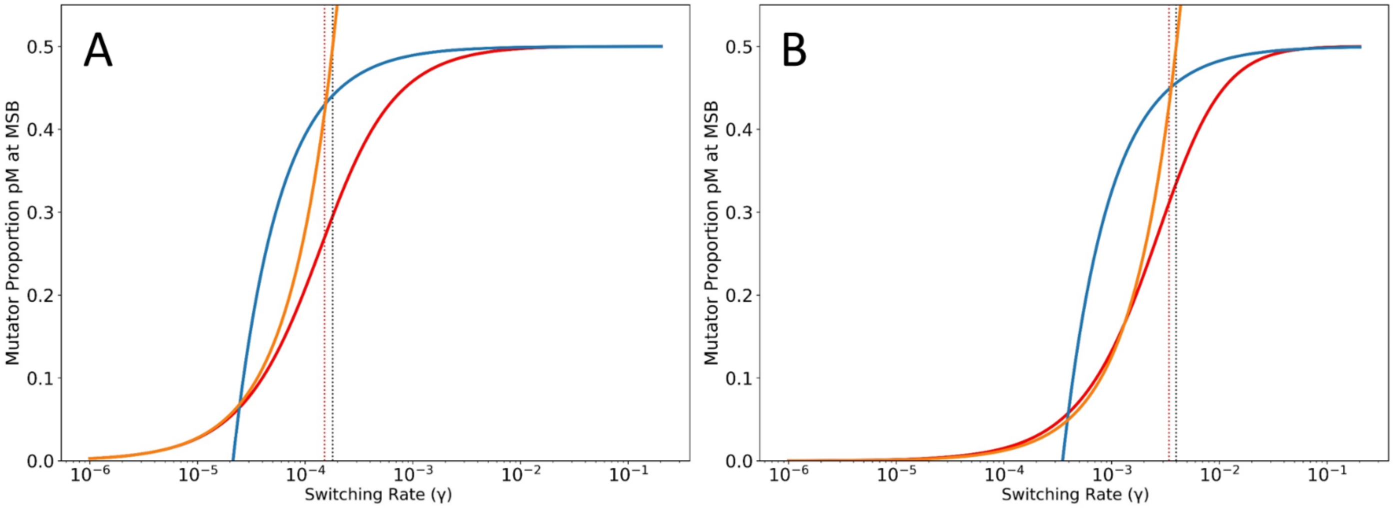


**Figure A1. Frequency of mutators $p_M$ at the mutation-selection balance.** Comparison of the mutant frequency $p_M$ from the deterministic model (red line; Eqs. 1-3) with the approximations $p_M^-$ (orange line; eq. A6) and $p_M^+$ (blue line; eq. A7). Although the exact transition point between the two approximations is given by the red dotted line (Eq. A10), we find that in practice the much simpler expression represented by the black dotted line (Eq. A11) provides a good approximation for the transition point. (A) $\boldsymbol{U = 0.00004}$; $\boldsymbol{s = 0.03}$; and $\boldsymbol{\tau = 10}$. Proportion of variance explained by the approximation: $\boldsymbol{R^2 = 0.966}$. (B) $\boldsymbol{U = 0.00004}$; $\boldsymbol{s = 0.03}$; and $\boldsymbol{\tau = 200}$. Proportion of variance explained by the approximation: $\boldsymbol{R^2 = 0.987}$.

### Frequency of Mutants, $p_S$

The number of deleterious alleles per individual at the MSB, with the same mutation rate in the entire population, follows a Poisson distribution with mean $U/_S$ (1). Therefore, the frequency of wild type (frequency of individuals with no mutant alleles) is

$$p_S^- = e^{-\frac{U}{s}}. \quad \text{(A12)}$$

With two mutation rate phenotypes (mutator and non-mutator), and when the phenotype switching rate is low, mutators are quickly purged from the population by selection, and the frequency of mutants $p_S$ tends to $p_S^-$. On the other hand, when the switching rate is very high and tends to 0.5, the population mean mutation rate tends to the average between the mutator and the non-mutator mutation rates, and we have

$$p_S^+ = e^{-\frac{U+\tau U}{2s}}. \quad \text{(A13)}$$

Between the two extremes, $p_S^-$ and $p_S^+$, the frequency of mutants $1 - p_S$ will increase with the switching rate. The rate of increase will be proportional to the population mean mutation rate. We find that a logistic function is a good approximation for the mutant frequency as a function of the switching rate, $p_S(\gamma)$. This logistic approximation, denoted $\tilde{p}_S$, is characterized by a rate $\frac{2}{U(\tau+1)}$, equal to the inverse of the average mutation rate when $\gamma = 0.5$, and left and right asymptotes at $p_S^-$ and $p_S^+$, respectively, namely

$$\tilde{p}_S = 1 - \frac{p_S^- \cdot p_S^+}{p_S^- + (p_S^+ - p_S^-) \cdot e^{-\frac{2\gamma}{U(\tau+1)}}} \quad \text{(A14)}$$

This approximation provides a good fit (**Figure A2**) and an intuitive understanding of $p_S$ as a function of the switching rate.

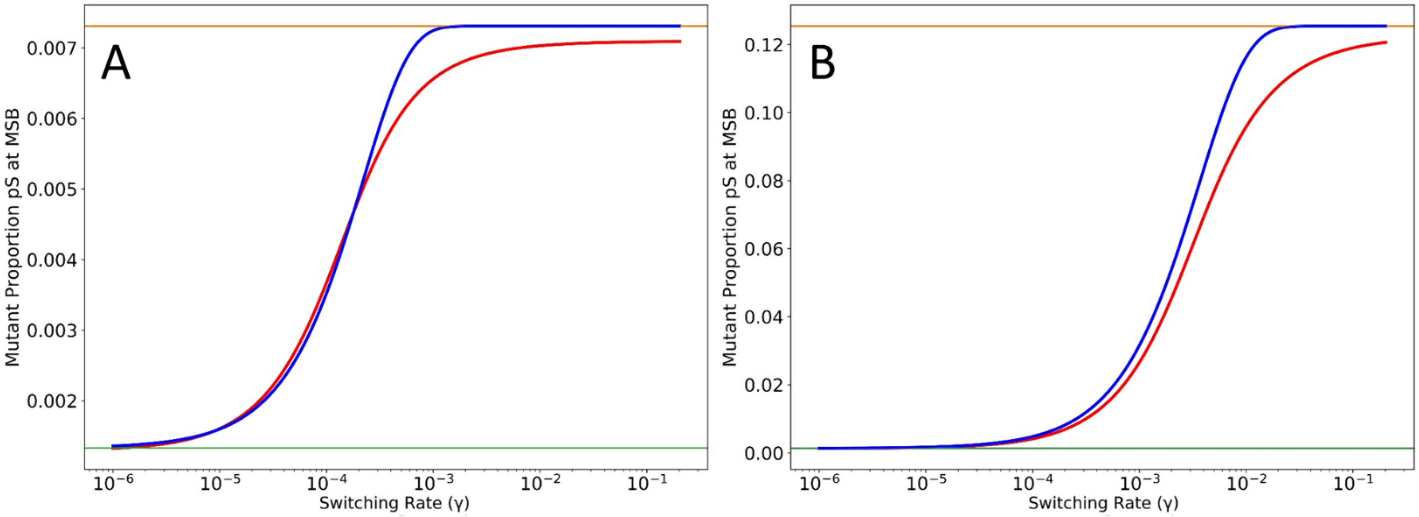


**Figure A2. Frequency of mutants $p_S$ at the mutation-selection balance.** Comparison of the mutant frequency $p_S$ from the deterministic model (red line; Eqs. 1-3) with the logistic approximation $\tilde{p}_S$ (blue line; eq. A14). The asymptotes for low and high switching rates, $p_S^-$ and $p_S^+$, are shown in green and orange respectively. (A) $U = 0.00004$; $s = 0.03$; and $\tau = 10$. Proportion of variance explained by the approximation: $R^2 = 0.996$. (B) $U = 0.00004$; $s = 0.03$; and $\tau = 200$. Proportion of variance explained by the approximation: $R^2 = 0.994$.

**Ratio of mutant mutator to wild type mutator, $p_R$**

We define $p_R = m_1/m_0$, that is, the ratio of non-mutator mutant frequency, *m1*, to non-mutator wildtype frequency, *m0*. We find that a logistic function with coefficient $\frac{\log(\tau)}{s}$ provides a good approximation and an interpretable understanding of $p_R$ as a function of the switching rate. As for the asymptotes: When the switching rate is very low, most mutators will be purged from the population before switching back to non-mutator. Therefore, the proportion of mutants among the non-mutator subpopulation will be independent of the distribution among the mutator subpopulation. Thus, $m_0$ will be close to 1 and $m_1$ will be close to 1-$p_S$, which is $1 - p_S^-$ when $\gamma$ is low. Hence the left asymptote of the logistic function will equal $1 - p_S^-$.
When the switching rate is high, close to 0.5, a non-mutator individual has a 50% chance of being the offspring of a mutator, and so we have $\frac{m_1}{m_0} = \frac{M_1}{M_0} = \frac{0.5\,(1-p_S)}{0.5\,p_S} \approx 1 - p_S^+$, since $1 - p_S^+$ will always be very small.
Therefore, $p_R$ will tend to $1 - p_S^+$ as the switching rate approaches 0.5.
Therefore, we approximate the ratio of mutant to wild type among non-mutators, $p_R$, with

$$\tilde{p}_R = \frac{p_S^- \cdot p_S^+}{p_S^- + (p_S^+ - p_S^-) \cdot e^{-\log(\tau)\frac{\gamma}{s}}}. \qquad \text{(A15)}$$

This approximation provides a good fit (**Figure A3**) and an interpretable understanding of $p_R$ as a function of the switching rate.

In **Figure A4,** we show the agreement between the approximations for $m_0$, $m_1$, $M_0$ and $M_1$ obtained from Eqs. A5B, A6, A7, A11, A14 and A15 and the results of the numerical iterations of Eqs. (1-3).

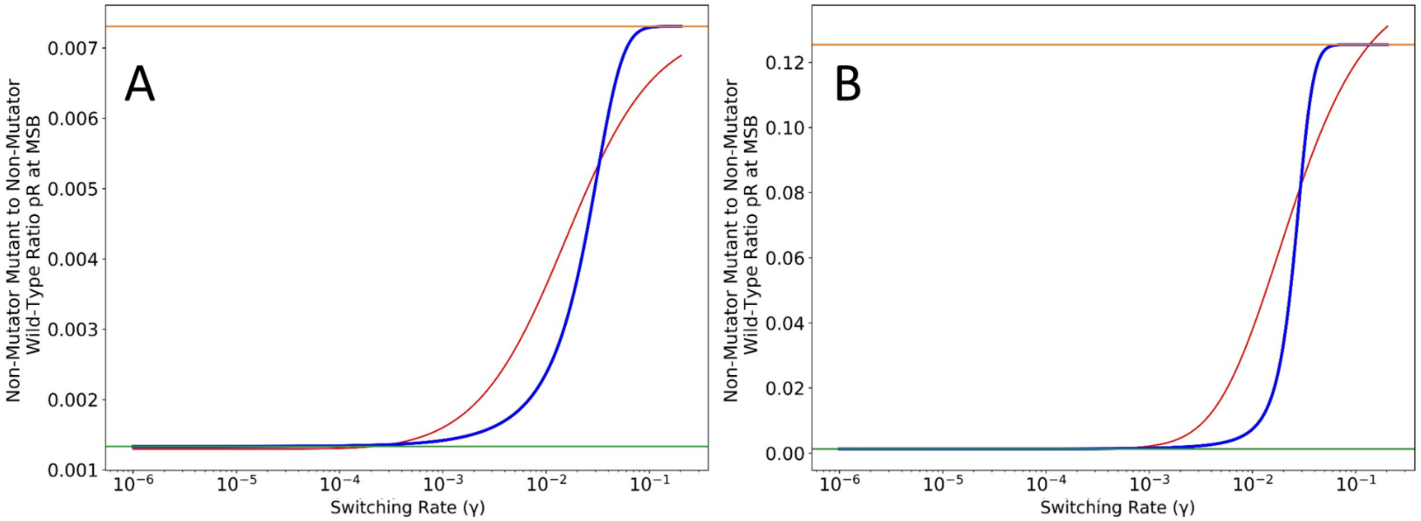


**Figure A3. Ratio $p_R$ at the mutation-selection balance.** Comparison of $m_1/m_0$ ratio $p_R$ from the deterministic model (red line; Eqs. 1-3) with the logistic approximation $\tilde{p}_R$ (blue line; eq. A15). The asymptotes for low and high switching rates, $p_S^-$ and $p_S^+$, are shown in green and orange respectively. (A) $U = 0.00004$; $s = 0.03$; and $\tau = 10$. Proportion of variance explained by the approximation: $R^2 = 0.943$. (B) $U = 0.00004$; $s = 0.03$; and $\tau = 200$. Proportion of variance explained by the approximation: $R^2 = 0.934$.

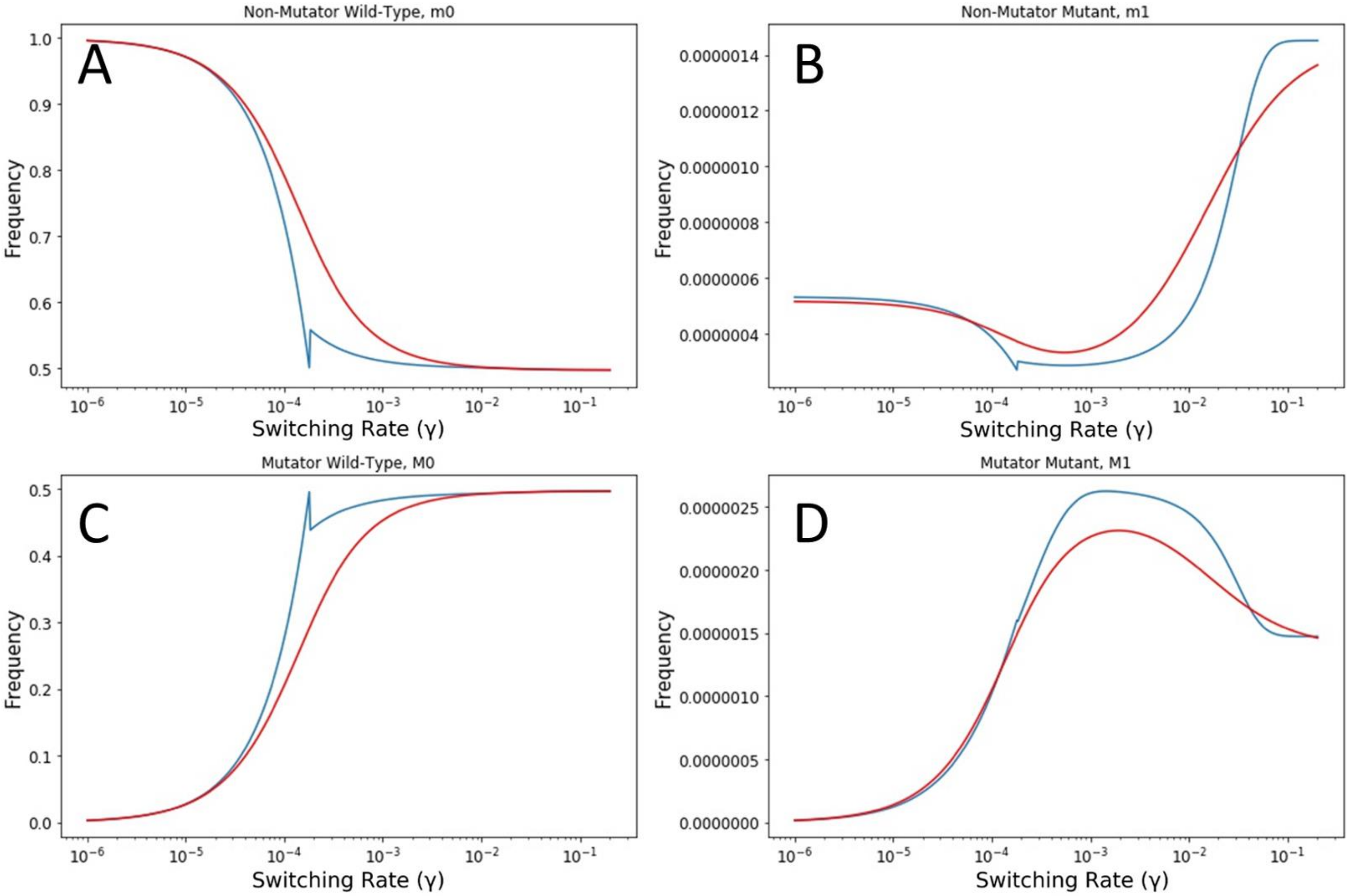


**Figure A4. Frequencies *m₀*, *m₁*, *M₀*, and *M₁*.** Comparison of the four frequencies ≈ from the deterministic model (red line; eqs. 1-3) with their approximations (blue; eqs. A5b). Parameters: $U = 0.00004$; $s = 0.03$; and $\tau = 10$. Proportion of variance explained by the approximations for $m_0$, $m_1$, $M_0$, and $M_1$: $R^2_{m0} = 0.967$; $R^2_{m1} = 0.935$; $R^2_{M0} = 0.966$ ; $R^2_{M1} = 0.989$, respectively.

### Fixation of an Adaptive Mutant

According to Eshel (4), in a large population with weak selection, the fixation probability $\rho_F$ of a double mutant once it appears in a single copy depends only on the selection coefficient $s$ and the double mutant advantage $H$, namely

$$\tilde{\rho}_F = \frac{2sH}{1+sH} \quad \text{(A16a)}$$

$$\approx 2sH, \quad \text{(A16b)}$$

where eq. A16b applies when $sH$ is much smaller than 1.

**Figure A5** shows a comparison of the two approximations in eq. A16 to results from stochastic simulations obtained by counting the number of fixation events after the appearance of an adaptive genotype**.**

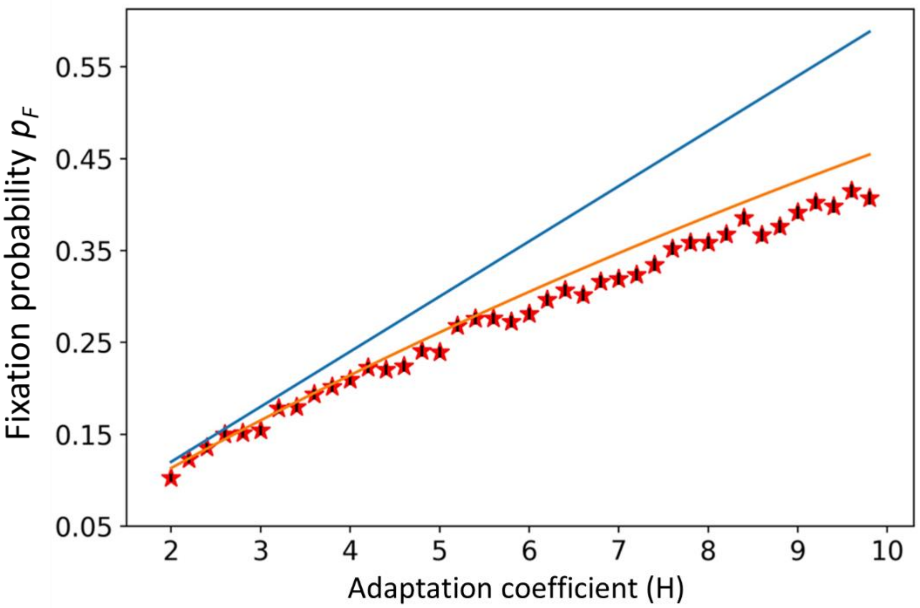


**Figure A5. Fixation probability of a rare beneficial genotype.** Red stars represent the frequency of fixations in *n*=5000 events of adaptive genotype appearance. Error bars (too small to see) show the estimated error $\sqrt{\boldsymbol{p_F(1-p_F)/n}}$. The analytic approximation in Eq. A16a (orange) explains $\boldsymbol{R^2} = \boldsymbol{0.9968}$ of the variance in the simulation results, while the approximation Eq. A16b (blue) explains $\boldsymbol{R^2} = \boldsymbol{0.9919}$. Here, *s = 0.03*.

We notice that Eq. A16b is independent from the switching rate $\gamma$. Hence, the adaptation rate will be proportional to the appearance rate (see **Figure A6** and **Figure 2**). We therefore consider the appearance rate as a proxy for the adaptation rate in the main text.

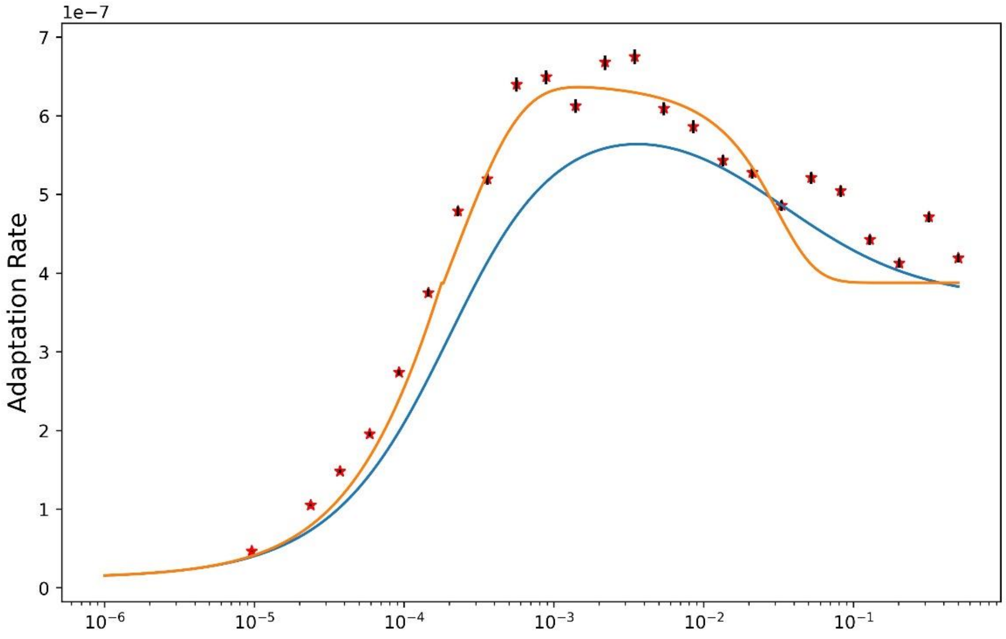


**Figure A6. The adaptation rate versus switching rate is proportional to the appearance rate.** This is because Eq. 16a is independent from the switching rate $\gamma$. Hence, in the main text, we focus on the appearance rate as a proxy for the adaptation rate. Stars show the adaptation rate on the two-peak landscape estimated from simulations (the stochastic model with genetic drift). Lines show the approximation (Eq. 6) using MSB frequencies from an analytic approximation (orange; Appendix E, Eqs. A3-A16) or numerical analysis (blue; Eqs. 1-3 without drift). Parameters: mutation rate, $\mu = 4 \cdot 10^{-5}$; mutation rate increase, $\tau = 10$; selection coefficient, $s = 0.03$; adaptation coefficient, H=5, population size $N = 10^7$. Error bars represent the 90% CI calculated according to augmented MCMC approach presented in Appendix A.

### Appendix F. Adaptation-optimal switching rate

We found that the adaptation rate is maximized by intermediate switching. To find the range of such intermediate rates, remember that the appearance of double mutants is (Eq. 5)

$$q = \mu\, e^{-U} m_1 + \tau\mu\, e^{-\tau U} M_1. \quad \text{(A17)}$$

Since $U<1$, the effect of $M_1$ on $q_A$ is larger than the effect of $m_1$ (**Figure A7**).

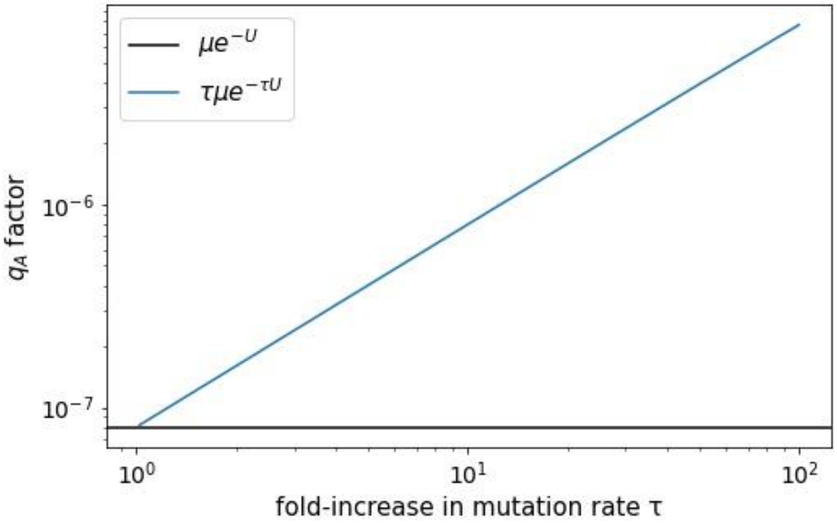

**Figure A7: Comparison of $\boldsymbol{\mu\, e^{-U}}$ and $\boldsymbol{\tau\mu\, e^{-U}}$ versus $\boldsymbol{\tau}$.** We notice that $\mu\, e^{-U}$is negligible with regards to $\tau\mu\, e^{-\tau U}$ for almost all of the considered values of $\tau$, the fold-increase in mutator mutation rate.

That is, the adaptation rate is maximized by switching rates that maximize $M_1$. From Eq. A5B, we have $M_1 = p_S - \frac{p_R}{1+p_R}(1 - p_M)$. If we assume $p_R$ is small ($m_1 \ll m_0$) then $\frac{p_R}{1+p_R} \approx p_R$. Hence, we have

$$\frac{M_1}{p_S} \approx 1 - \frac{p_R(1 - p_M)}{p_S}. \quad \text{(A18)}$$

As $\gamma$ tends to 0, $p_R$ and $p_S$ tend to $p_S^-$ and $1 - p_M$ tends to 1 (see *Appendix E*). Hence $M_1/p_S$ tends to 0. When $\gamma$ tends to 0.5, $p_R$ and $p_S$ tend to $p_S^+$ and $1 - p_M$ tends to 0.5.

Moreover, both functions, $p_S$ (Eq. A14) and $p_R$ (Eq. A15) are logistic functions, characterized by an initial low derivative with respect to $\gamma$, followed by a large derivative before finally plateauing at an asymptotic value (i.e., derivative approaches zero). $p_S$ and $p_R$ share asymptotic values: the lower asymptote, $p_S^-$, is the frequency of wild type (individuals with no mutations) if the whole population was non-mutator. The upper asymptote, $p_S^+$, is the frequency of wild type (individuals with no mutations) if the population was composed in equal parts of mutators and non-mutators, which occurs at a random switching rate. For both functions, the maximal derivative with respect to $\gamma$ will be reached at

$$argmax(p_S{}'(\gamma)) = \frac{\log(p_S^-/(p_S^- + p_S^+))}{2/U(\tau - 1)}, \quad \text{(A19A)}$$

for $p_S$ and

$$argmax(p_R'(\gamma)) = \frac{\log(p_S^-/(p_S^- + p_S^+))}{\log(\tau)\,/s}, \quad \text{(A19B)}$$

For $p_R$.

Taking $\log(p_S^-/(p_S^- + p_S^+)) \sim 1$, we obtain $argmax(p_S{}'(\gamma)) \sim \tau(U - 1)/2$ for $p_S$ and $argmax(p_R'(\gamma)) \sim s/\log(\tau)$ for $p_R$. Also $p_M$, a piecewise function of $p_M^-$ (Eq. A6) and $p_M^+$ (Eq. A7), is characterized by an initially low derivative that increases with increasing $\gamma$, before switching to a function with a lower derivative at the transition point $\tau(U - 1)/2$.

Therefore, $M_1/p_S$ will grow until reaching its maximum with maximal derivative reached at $(\tau - 1)U/2$. Then, as $p_R$ reaches its largest derivative, $p_R$ will then decrease with maximal negative derivative reached at $s/\log(\tau)$.

Hence, the adaptation-optimal switching rate $\gamma^*$ will be such that $(\tau - 1)U/2 < \gamma^* < s/\log(\tau)$.

The upper and lower bounds for the adaptation-optimal switching rate $\gamma^*$ versus the relevant parameters (the fold-increase in the mutator mutation rate $\tau$, and the non-mutator mutation rate $U$ for the lower bound; the fold-increase in the mutator mutation rate $\tau$, and the selection coefficient $s$ for the upper bound) are plotted in **Figure A8**.

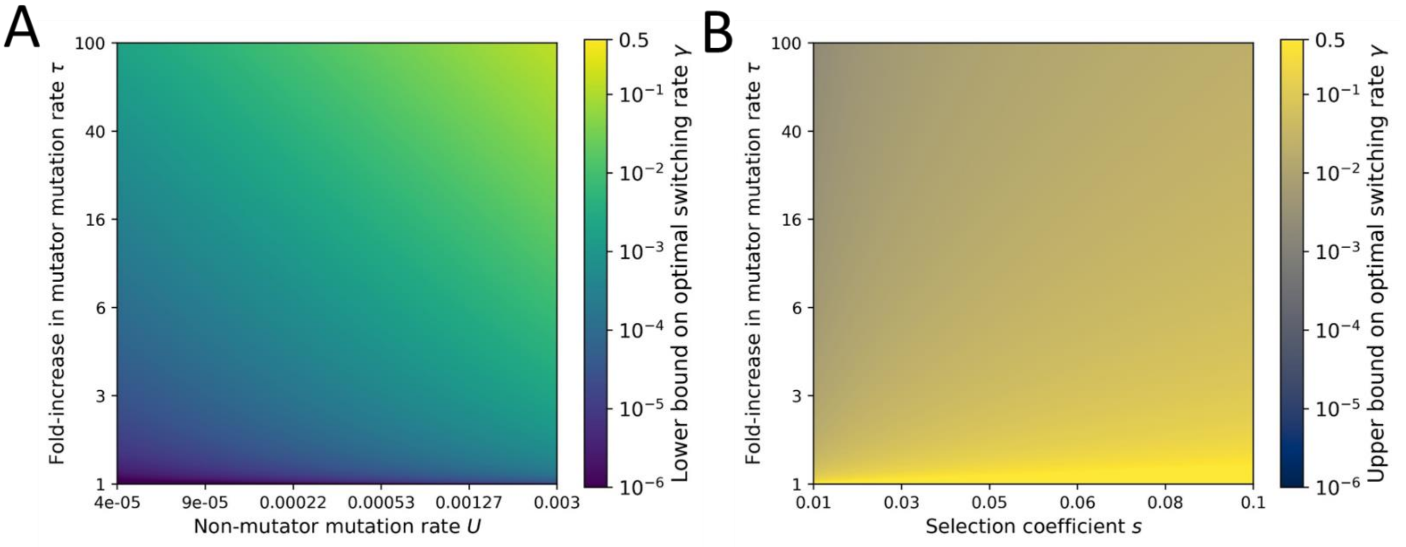


**Figure A8: (A) Lower bound on adaptation-optimal switching rate $\gamma^*$.** The lower bound is given by $(\tau - 1)U/2$. **(B) Upper bound on adaptation-optimal switching rate $\gamma^*$.** The upper bound is given by $s/\log(\tau)$. We notice that the adaptation-optimal switching rate is almost always higher than expected by genetic switching rates (estimated at about $10^{-6}$, see (5)) and lower than expected by a random switching rate (at 0.5).

### <u>Appendix G. Gradual transition between mutator and non-mutator phenotypes</u>

At very high switching rates, the effect of selection against the mutator on its frequency is negligible. Indeed, the distribution of the mutation rate phenotypes is regenerated at every generation. Hence, the equilibrium frequencies of the non-mutator, transient, and mutator phenotypes are given by the stationary distribution of the transition matrix that describes the transition probabilities between the three mutation rate phenotypes (see Eq. 4). Since the transition probabilities for the mutator and non-mutator phenotypes are symmetric, the stationary frequencies of these two phenotypes will be the same. The stationary frequency of the transient mutation rate phenotype will be non-zero; hence, the stationary frequency of the mutator phenotype at high switching rates will be less than 0.5. Since the other two mutation rate phenotypes have a lower mutation rate, the rate of adaptation will also be lower, as seen in **Figure 2**. The frequencies of the three mutation rate phenotypes as a function of the switching rate are shown in **Figure A10**.

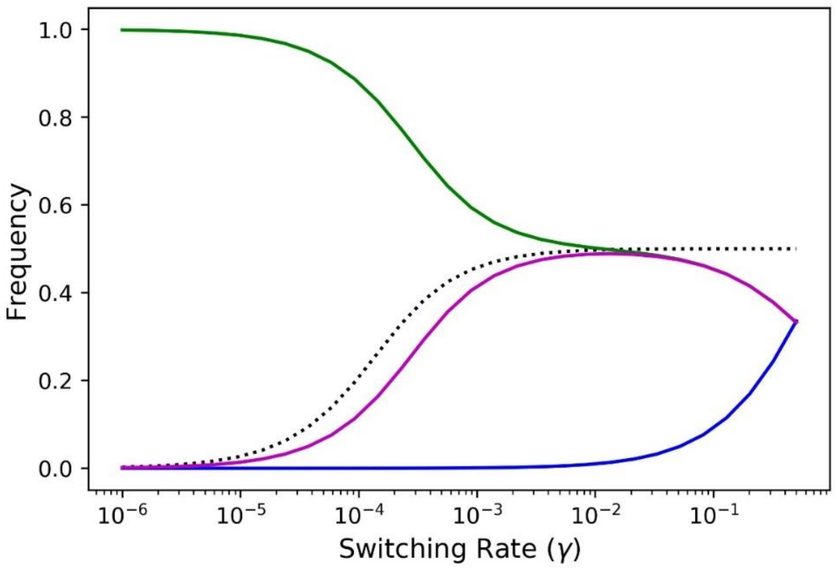


**Figure A9. Frequencies at MSB of the three mutation rate phenotypes in a model with gradual transition between mutator and non-mutator phenotypes.** Frequencies are given by the eigenvector corresponding to the leading eigenvalue of the mutation-selection matrix (see Table 1 and Eq.4). Green line for the frequency of the non-mutator phenotype; purple line for the frequency of the mutator phenotype; dark blue line for the frequency of the phenotype with a transient mutation rate equal to the geometric mean of the non-mutator and mutator mutation rate; dotted line for the frequency of the mutator phenotype from the two-phenotype model. We observe that the frequency of the mutator phenotype is higher in the two-phenotype model for very high switching rates, but similar for intermediate switching rates, which explains the higher ratio between the adaptation rates achieved for epigenetic switching rates and the adaptation rates achieved for random switching rates in the model with gradual transition between mutator and non-mutator phenotypes. Parameters: $\boldsymbol{U = 4 \cdot 10^{-5}}$; $\boldsymbol{s = 0.03}$; $\boldsymbol{\tau = 10}$; $\boldsymbol{\delta = 0.99}$.

### Appendix H. Effective selection coefficient in NK and empirical landscapes

The effective selection coefficient *s* is defined as the average effect of deleterious mutations on the fitness, such that the fitness of a mutant is (1-s). Therefore, the effective selection coefficient *s* for the three considered NK landscapes and the *Aspergillus niger* landscape is

$$s = \frac{w_{wt} - \overline{w}_{mt}}{w_{wt}}, \tag{A20}$$

where $s$ is the effective selection coefficient, $w_{wt}$ is the fitness of the wildtype genotype, and $\overline{w}_{mt}$ is the average fitness of single mutants (6).

In practice, we found high variance of single mutant fitness values, such that some mutants are much less frequent than others at the MSB, and so their fitness can be neglected in the estimation of s. Thus, we estimated *s* (Eq. A20) by substituting $\overline{w}_{mt}$ with the fitness of the fittest single mutant (which is also the most frequent mutant), with the average of the fitness of the two fittest single mutants, with the average of the fitness of the three fittest mutants, etc. The effective selection coefficients *s* were calculated considering the *x* "neighbours" for *x* between 1 and 8. For each value, the Euclidean distance between the approximation and the deterministic model was computed and the *s* value resulting in the lowest Euclidean distance was chosen. The results for the NK landscapes are shown in **Figure A10**. In the least rugged landscape (*k=1*), the best approximation was obtained by considering almost all single mutants of the wildtype (**Fig. A10A**). In the most rugged landscape ($k$=5), the best approximation was obtained by considering only the fittest of the single mutants (**Fig. A10C**). Results for the empirical fitness landscape are in **Figure A11**.

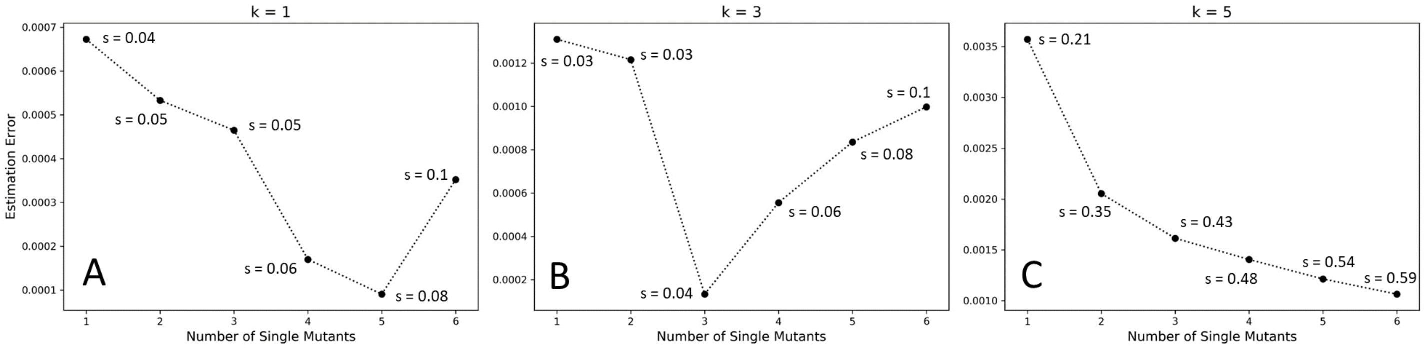


**Figure A10. Effective selection coefficient in NK fitness landscapes**. The effective selection coefficient *s* was estimated as the average of the most fit mutants of the wildtype (x-axis). The estimation error, i.e. Euclidean distance, between the estimation and rate of appearance calculated with $m_0$, $m_1$, $M_0$, and $M_1$ values from the deterministic model (y-axis) was computed, and the *s* with the smallest estimation error was chosen. Parameters: $\boldsymbol{\mu = 10^{-6}}$; $\boldsymbol{\tau = 10}$, s = 0.04, N = 100,000.

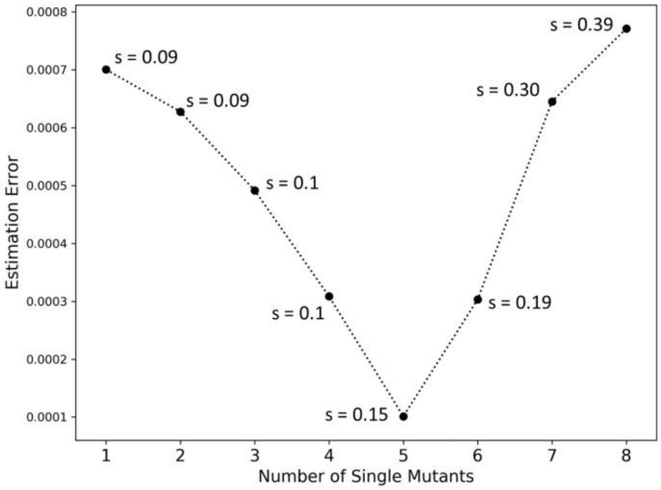


**Figure A11. Effective selection coefficient in *Aspergillus niger* empirical fitness landscapes.** The effective selection coefficient *s* was estimated as the average of the most fit mutants of the wildtype (x-axis). The estimation error, i.e. Euclidean distance, between the estimation and rate of appearance calculated with $m_0$, $m_1$, $M_0$, and $M_1$ values from the deterministic model (y-axis) was computed, and the *s* with the smallest estimation error was chosen. Parameters: $\boldsymbol{\mu = 10^{-6}}$; $\boldsymbol{\tau = 10}$, s = 0.04, N = 100,000.

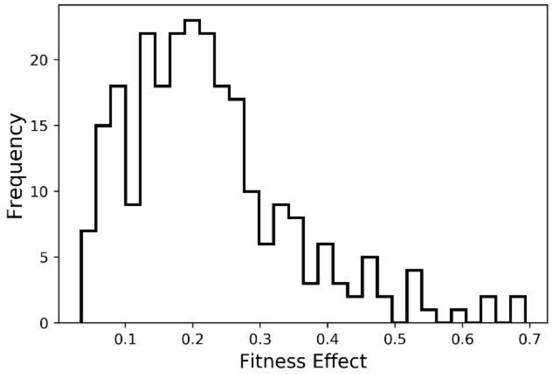


**Figure A12. Distribution of fitness effects in the *Aspergillus niger* empirical landscape.** A histogram of the mean difference between each genotype and its single mutants.

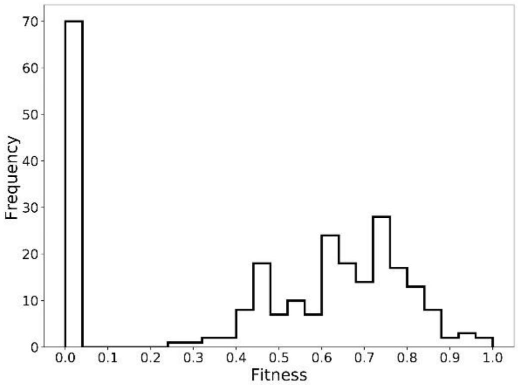


**Figure A13. Distribution of fitness values in the *Aspergillus niger* empirical landscape.** Fitness values are growth rates relative to the wildtype.

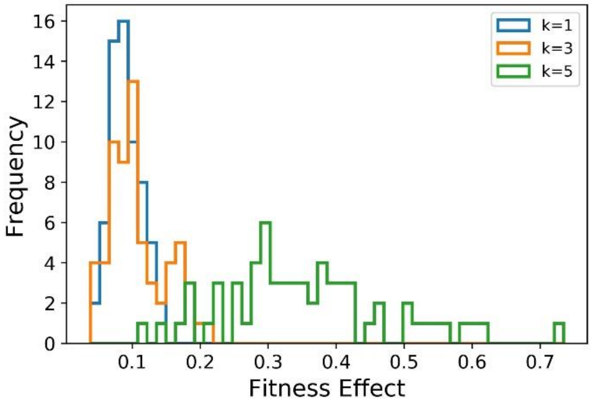


**Figure A14. Distribution of fitness effects in the NK landscapes.** A histogram of the mean difference between each genotype and its single mutants for different values of the ruggedness parameter *k*. As expected, as *k* increases, the distribution is wider.

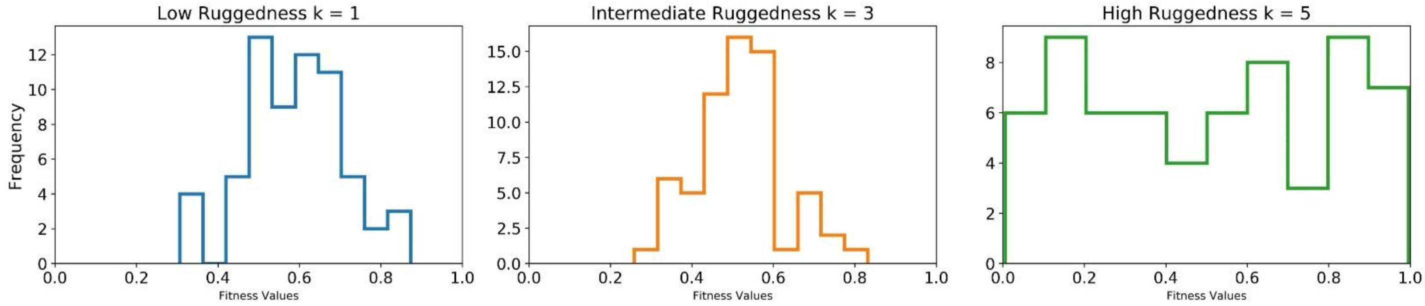


**Figure A15. Distribution of fitness values in the NK landscapes.** A histogram of the fitness values of all genotypes for different values of the ruggedness parameter, *k*. As expected, as *k* increases, the distribution is wider. Blue indicates $\boldsymbol{k = 1}$. Orange indicates $\boldsymbol{k = 3}$. Green indicates $\boldsymbol{k = 5}$.

| k | Wildtype genotype | Fitness of wildtype genotype | Mean fitness of single mutants | Minimal fitness difference | Maximal fitness difference | Minimal fitness of single mutants | Global maximum genotype | # local maxima in landscape |
|---|---|---|---|---|---|---|---|---|
| 1 | 111100 | 0.71 | 0.64 | 0.03 | 0.17 | 0.54 | 100111 | 1 |
| 3 | 100111 | 0.57 | 0.52 | 0.01 | 0.11 | 0.47 | 111010 | 5 |
| 5 | 100101 | 0.93 | 0.38 | 0.2 | 0.81 | 0.12 | 001110 | 7 |

**Table A2. Fitness of wildtype genotype and its single mutants in the NK landscapes.**

#### A) *Two-Peak* Landscape

| | Analytic Approximation | Numerical Iteration | Simulations |
|---|---|---|---|
| Analytic Approximation | 1 | 0.96 | 0.96 |
| Numerical Iteration | - | 1 | 0.93 |
| Simulations | - | - | 1 |

#### B) *NK = 3* Landscape

| | Analytic Approximation | Numerical Iteration | Simulations |
|---|---|---|---|
| Analytic Approximation | 1 | 0.98 | 0.94 |
| Numerical Iteration | - | 1 | 0.98 |
| Simulations | - | - | 1 |

#### C) *Aspergillus niger* Empirical Landscape

| | Analytic Approximation | Numerical Iteration | Simulations |
|---|---|---|---|
| Analytic Approximation | 1 | 0.99 | 0.93 |
| Numerical Iteration | - | 1 | 0.99 |
| Simulations | - | - | 1 |

**Table A3. Coefficients of determination $R^2$ between deterministic, stochastic, and analytical approximation of the model.**

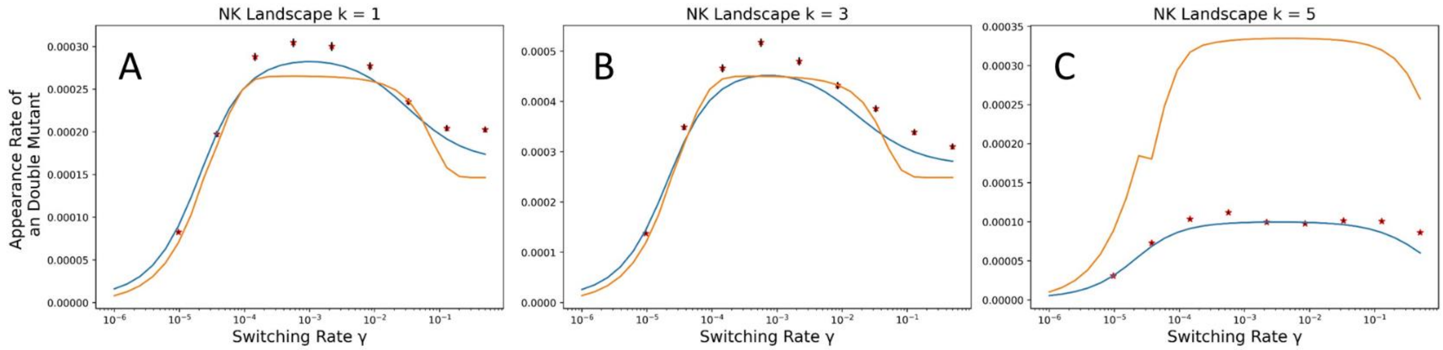


**Figure A16. Appearance rate of double mutants maximized by intermediate switching rates in the NK fitness landscape.** Moreover, we observe a good fit between the rate of appearance computed with MSB frequencies from the analytical approximation (Eqs. A5b, 5) and the appearance rate with MSB frequencies from the eigenvector corresponding to the leading eigenvalue of the mutation-selection matrix (see Table 1 and Eq.3). $\boldsymbol{R^2}$ represent the correlations between the rate of appearance calculated from the approximations presented in the *Appendix E* and the rate of appearance calculated from the numerical iterations of the model. Parameters: $\boldsymbol{U = 10^{-6}}$; $\boldsymbol{\tau}$ = 10; (A): s = 0.0630; (B): s = 0.0443; (C): s = 0.5955; (A), (B): N = 100000; (C) N = 1000000. Error bars represent 90% CI calculated with the augmented MCMC method (see Appendix A).
